# Event-related modulation of alpha rhythm explains the auditory P300 evoked response in EEG

**DOI:** 10.1101/2023.02.20.529191

**Authors:** A.A. Studenova, C. Forster, D.A. Engemann, T. Hensch, C. Sander, N. Mauche, U. Hegerl, M. Loeffler, A. Villringer, V.V. Nikulin

## Abstract

Evoked responses and oscillations represent two major electrophysiological phenomena in the human brain yet the link between them remains rather obscure. Here we show how most frequently studied EEG signals: the P300-evoked response and alpha oscillations (8–12 Hz) can be linked with the baseline-shift mechanism. This mechanism states that oscillations generate evoked responses if oscillations have a non-zero mean and their amplitude is modulated by the stimulus. Therefore, the following predictions should hold: 1) the temporal evolution of P300 and alpha amplitude is similar, 2) spatial localisations of the P300 and alpha amplitude modulation overlap, 3) oscillations are non-zero mean, 4) P300 and alpha amplitude correlate with cognitive scores in a similar fashion. To validate these predictions, we analysed the data set of elderly participants (N=2230, 60–82 years old), using a) resting-state EEG recordings to quantify the mean of oscillations, b) the event-related data, to extract parameters of P300 and alpha rhythm amplitude envelope. We showed that P300 is indeed linked to alpha rhythm, according to all four predictions. Our results provide an unifying view on the interdependency of evoked responses and neuronal oscillations and suggest that P300, at least partly, is generated by the modulation of alpha oscillations.

## Introduction

P300 is one of the most extensively investigated evoked responses (ER) in electroencephalography (EEG) and magnetoencephalography (MEG). Over the years, P300 has been hypothesised to reflect a variety of functions, such as priming, cognitive processing, memory storage, context updating, resource allocation, etc. (Polich et al., 1995, Polich, 2003, Verleger, 2020), and there is an ongoing effort to understand its functions further through such constructs as information, expectancy, and capacity (Verleger, 2020). Usually, P300 is assessed with the oddball paradigm (auditory or visual), where participants have a task to detect a target (or rare, or deviant) stimulus in a train of standard (or frequent, or non-target) stimuli (Luck, 2014). Additionally, it is usual to speak about the P300 complex, involving the earlier frontal component P3a and the later parietal component P3b (Linden, 2005). Being aware of this forking terminology, in the following, we refer to P300 as the ER that occurs after the target stimulus and is different compared to the ER to the standard stimulus. Adding to the complexity of P300, the exact mechanism of P300 generation remains rather unknown (Fell et al., 2004, Hanslmayr et al., 2007, Daly et al., 2009, Rawls et al., 2020). In the present study, we investigate a possibility that P300 might be to some extent generated through a baseline-shift mechanism (BSM, Nikulin et al., 2007, Mazaheri et al., 2008, Iemi et al., 2019, Studenova et al., 2022).

Apart from P300, the oddball target stimulus concurrently causes the attenuation of the alpha rhythm amplitude (8–12 Hz). The simultaneity of P300 and alpha rhythm modulation has been observed in numerous earlier and more recent studies, and in Supplementary material we offer a short overview of these findings (Table S1). We found 38 studies that presented results for a concomitant occurrence of P300 and alpha power (or amplitude). In 17 studies using EEG, results indicated an overlap in cortical regions of P300 and alpha amplitude decrease, as well as a similar time windows of their occurrence (Peng et al., 2012, Chen et al., 2013, Dong et al., 2015, Shou et al., 2015, Tang et al., 2015, Wu et al., 2015, Fabi et al., 2017, López-Caneda et al., 2017, Vilà-Balló et al., 2017, Fabi et al., 2018, Michelini et al., 2018, Román-López et al., 2019, Kao et al., 2020, Yu et al., 2020, Zhang et al., 2020, Nikolin et al., 2021, Paolicelli et al., 2021). Similar observations were made using MEG (Ishii et al., 2009). Yordanova et al. (2001) and 14 more studies found similarities in location but not in the peak latencies (Kolev et al., 2001, Kamarajan et al., 2006, Digiacomo et al., 2008, Krämer et al., 2011, Barutchu et al., 2013, Deiber et al., 2013, Kayser et al., 2013, Zarka et al., 2014, Deiber et al., 2015, Leroy et al., 2017, Liu et al., 2019, Martel et al., 2019, Faro et al., 2020, Espenhahn et al., 2020). In only a few studies, alpha modulation did not appear at all (Kamarajan et al., 2004, Delval et al., 2018) or the relationship between alpha oscillations and ER was not supported by cross-condition comparison (Cooper et al., 2008, Lee et al., 2017, Tamura et al., 2016). In general, we acknowledge that due to different ways of presenting results, sometimes it was difficult to tell whether the peak of P300 and the attenuation peak in the alpha amplitude correspond to each other. Nevertheless, the vast majority of studies confirmed the simultaneous occurrence of P300 and alpha amplitude decrease in several experimental paradigms, which in turn served as a basis for further investigation carried out in the present study.

The simultaneous presence of P300 and alpha amplitude modulation in the poststimulus window indicates that P300 can be partially generated through BSM (Nikulin et al., 2007, Mazaheri et al., 2008, Iemi et al., 2019, Studenova et al., 2022). Previous research investigated whether the origin of P300 is due to an additive mechanism (Fell et al., 2004, Wan et al., 2009, Herrmann et al., 2014) or a phase-reset mechanism (Fell et al., 2004, Daly et al., 2009, Wan et al., 2009, but Popp et al., 2019), and the evidence for these mechanisms is far from converging. However, P300 has not yet been assessed with respect to BSM. In general theory, BSM links evoked activity and spontaneous oscillatory activity, stating that if oscillations are modulated by the stimulus presentation, this modulation will be mirrored in the low-frequency signal if oscillations have a non-zero mean (see Figure 1). In other words, the amplitude modulation of the oscillatory process affects the mean as well, which in turn leads to the deflection in the spectral range of modulation activity (with the frequency of modulation lying in a considerably lower range than the carrier frequency of oscillations themselves; for instance, if the oscillations’ frequency range is 8–12 Hz, the modulation’s frequency range is 0–3 Hz). In practice, when integrated over several periods, oscillations with a non-zero mean will show an average value different from zero that will scale with the amplitude of oscillations. Likewise, a non-zero mean implies that average values of the upper and lower half of the oscillatory cycle would be unequal. In Figure 1, negative-mean oscillations undergo a decrease in the amplitude in the poststimulus window and, according to BSM, the decrease in the amplitude of negative-mean oscillations creates an ER with a positive polarity. Here, oscillations are assumed to be ongoing, i.e., they are present before the stimulus onset. The polarity of the ER depends on the sign of the oscillatory mean and on the direction of modulation—an increase or decrease in the amplitude. The oscillatory mean of alpha oscillations has been shown to be present in biophysical model of alpha oscillations (Studenova et al., 2022) and several studies provided empirical evidence for the generation of ER through BSM in somatosensory (Nikulin et al., 2007) and visual (Mazaheri et al., 2008, Iemi et al., 2019) domain. Since P300 coincides with the stimulus-triggered decrease in the alpha amplitude, it is reasonable to assess the compliance of P300 with BSM. Therefore, we hypothesised that P300 generation can at least partially be explained by the amplitude modulation of alpha oscillations, and in the following, we offer a systematic investigation of this hypothesis.

**Figure 1.**
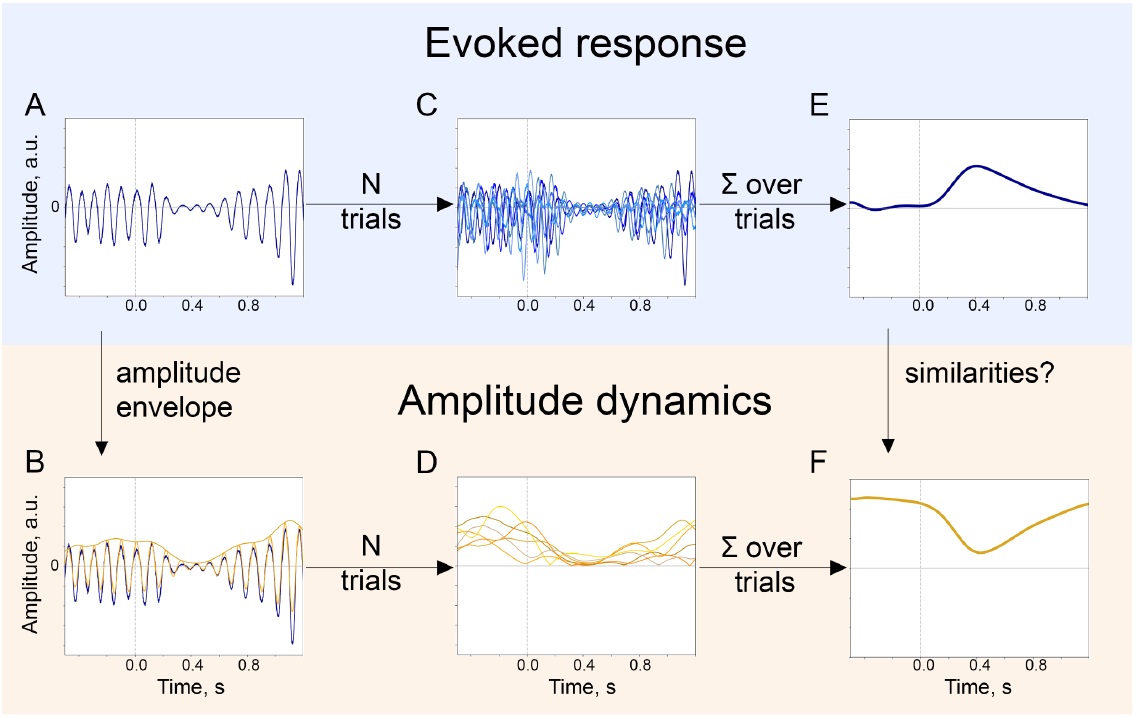
The baseline-shift mechanism (BSM) of evoked response (ER) generation. For a particular ER, probing the agreement with BSM would involve extracting both the ER and the oscillatory amplitude envelope. **A**. The single-trial broadband signal. **B**. The amplitude envelope of oscillations is extracted from a broadband signal of each trial. **C**. To get a high signal-to-noise ER, usually a few trials are acquired. Note that since oscillations have a negative mean, their attenuation would lead to the generation of an ER with a positive polarity (shown in E.). **D**. Similarly, for each trial, the amplitude envelope is extracted. **E**. Trials are averaged and, optionally, low-pass filtered to obtain an ER. **F**. Amplitude envelopes over trials are also averaged to obtain an estimate of the change in oscillatory amplitude in the poststimulus window. Here, we simulated the example of negative-mean oscillations giving rise to a positive-polarity ER.

Assessing the compliance of ER to BSM requires the following four prerequisites: 1) demonstrating the similarity in the temporal evolution of both signals—P300 and alpha amplitude envelope—over time in the poststimulus interval, 2) showing the similarity of spatial locations of the neuronal processes giving rise to P300 and to alpha amplitude decrease, 3) linking the direction of ER with the direction of alpha amplitude modulation through the sign of oscillatory mean, 4) establishing similarity of a relation of ER/oscillations with external variables, such as cognitive performance. In the following sections, we present comprehensive evidence for the association of P300 with alpha oscillations using a large EEG data set. In this data set, the experimental task was an auditory oddball paradigm. Participants would hear tones, one type of which—the target tone—would occur in only 12% of trials. Target tones elicit both P300 and the modulation of the alpha amplitude. Firstly, we show that in sensor space, the time courses of P300 and the alpha amplitude envelope are negatively correlated in the posterior region, and, in addition, the depth of alpha amplitude modulation correlates with the amplitude of P300. Secondly, we demonstrate that the increase in the low-frequency amplitude, that is P300, is pronounced over the posterior region, where at the same time the decrease in the alpha amplitude also occurs. Additionally, we perform source reconstruction to precise the location. Thirdly, by means of the baseline-shift index (BSI, Nikulin et al., 2010), we estimate oscillatory mean and establish that the sign of the mean is predictive of the P300-alpha relation. Finally, we evaluate the correlation between cognitive processes such as attention, memory, and executive function with P300 and alpha rhythm to confirm the relatedness of the two phenomena via behaviour.

## Results

### Temporal similarity between alpha amplitude envelope and P300

In line with the first prediction, average time courses of P300 and alpha amplitude envelope demonstrate an inverse relation—while P300 has a positive deflection, alpha rhythm amplitude is attenuated (Figure 2). The ER after the standard stimulus does not demonstrate the same strong relation (we will refer to ER after the standard stimulus as sER). To illustrate the relation even further, we filtered the ER in low frequency up to 3 Hz. Figure 2A on the left demonstrates the evolution of averaged time courses of ER at the Pz electrode, and Figure 2A on the right is the same but for the alpha amplitude envelope (see also Figure S3 for the whole-head time courses). This figure clearly shows a similarity in the temporal evolution for both types of signals. More specifically, within a window 200–400 ms after stimulus onset, P300 has a rising flank and alpha amplitude starts to decrease, and within a window 400–700 ms, both P300 and alpha amplitude have the largest magnitude (Figure 2B). To quantify this relation, we estimated the correlation between P300 and alpha amplitude envelope over averaged signals at every electrode. As predicted, the correlation for the target stimulus was significantly negative at posterior regions (Figure 2C, at Pz correlation is –0.86).

**Figure 2.**
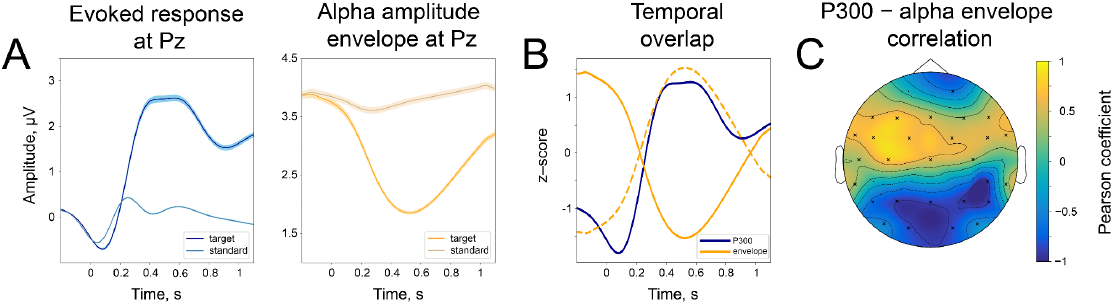
Temporal similarity between P300 and the alpha amplitude envelope. A. Left panel—time course of P300 at the Pz electrode elicited by the target stimulus and ER after a standard stimulus (sER) both averaged across participants. Right panel—alpha amplitude envelope at Pz electrode averaged across participants for target and standard stimulus. Shaded areas display the standard error of the mean. B. Temporal overlap in signals. The time courses of P300 and alpha amplitude display similarities in initial slope and peak latency. Amplitude values are z-scores to aid visual comparison. Dashed line—alpha amplitude envelope multiplied by –1. C. A correlation between P300 and alpha amplitude. For grand averages at each electrode, the correlation between P300 and alpha envelope was computed with the Pearson correlation coefficient. Electrodes marked with “x” had significant correlation coefficients. The p-value was set at the Bonferroni corrected value of 10^−4^. Note the positive correlation between the low-frequency signal and the alpha amplitude envelope over central sites. Due to the negative polarity of ER over the fronto-central sites, such correlation may still indicate a temporal relationship between the P300 process and oscillatory amplitude envelope dynamics (due to the use of a common average reference). However, it cannot be entirely excluded that additional lateralized response-related activity contributes to this positive correlation (Salisbury et al., 2001).

According to the baseline-shift mechanism, the change in the strength of the amplitude modulation should be mirrored in the change in P300 amplitude. Indeed, when we sorted alpha amplitude envelopes between participants into 5 bins according to the normalised change, the P300 amplitude followed the partition of alpha amplitudes. The normalised change was computed as 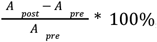, meaning that a value closer to –100 % corresponds to a strong drop in the poststimulus amplitude in comparison to prestimulus, while a value closer to 0 % corresponds to the absence of change in the amplitude, and a value larger than 0 % corresponds to the increase in the amplitude in the poststimulus window. The different alpha amplitude dynamics correlated with P300 amplitude, such that for participants with a stronger alpha amplitude modulation, the amplitude of P300 was higher than for participants with weak amplitude modulation (Figure 3A,B). As predicted by BSM, a smaller alpha amplitude modulation will generate an ER with a smaller amplitude. The total number of participants in each bin is 446. The t-test between the most extreme bins demonstrates a significant spatio-temporal cluster in the posterior region spanning electrodes CP6, P3, P4, P7, Pz, O1, O2, PO9 (Figure 3C). Here, t-values are negative, meaning that P300 that coincides with small alpha amplitude attenuation is significantly smaller in its amplitude than P300 that coincides with the largest alpha amplitude attenuation. The cluster within the earlier window (100–200 ms) over central regions (Figure 3C) possibly reflects the previously shown effect of prestimulus alpha amplitude on earlier ERs (Brandt et al., 1991, Babiloni et al., 2008) but may also be a manifestation of BSM. We tested this assumption for early ER, which in our auditory task was N100. We repeated the binning analysis for broadband data (0.1–45 Hz) and also observed a significant difference between two extreme bins around 100 ms over the central region (Figure S5A). However, if we filter the signal from 4 to 45 Hz (the range that includes the frequency of N100 but not low-frequency baseline shifts), these significant differences almost completely disappear (only electrode TP9 was significant; Figure S5B). It means that the difference in N100 amplitudes over frontal sites is driven by the baseline shift created by an unfolding alpha amplitude decrease. The significant difference at the TP9 electrode possibly reflects a genuine physiological effect of alpha rhythm amplitude on the excitability of a neuronal network and, as a consequence, on the amplitude of ER (as opposed to the baseline-shift mechanism, where the alpha rhythm doesn’t affect the amplitude of ER but creates an additional component of ER; Iemi et al. 2019).

**Figure 3.**
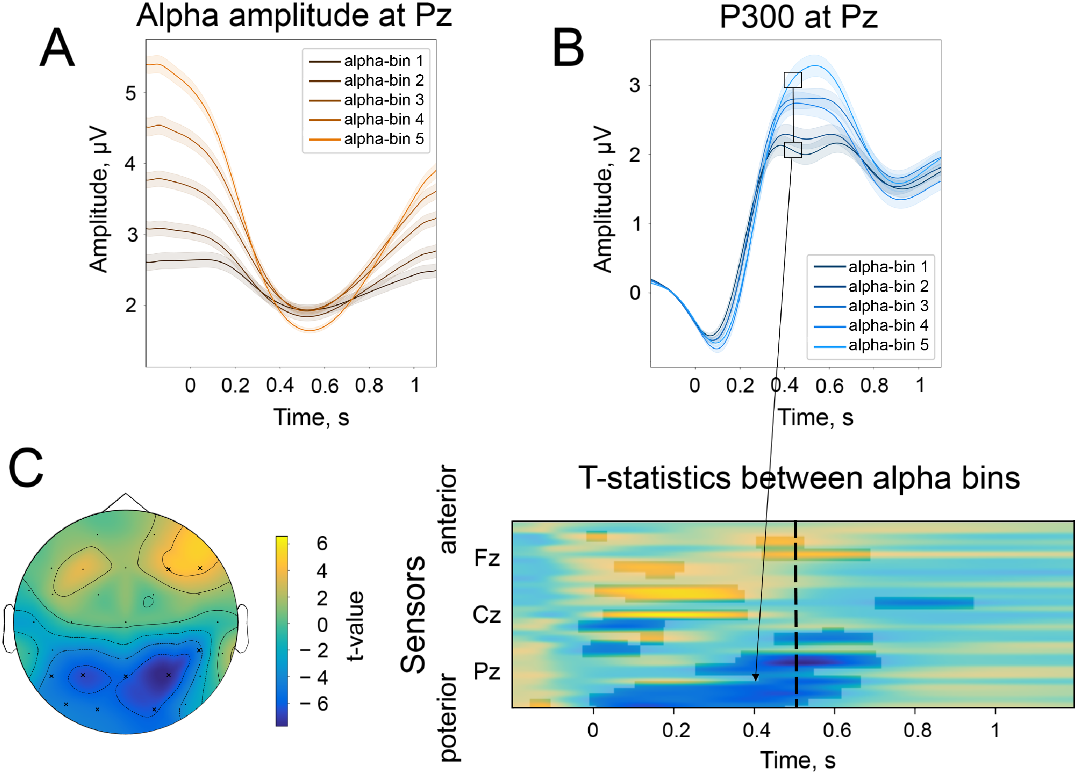
The difference in the strength of alpha amplitude modulation correlates with the difference in P300 amplitude. **A**. Alpha amplitude envelope sorted into 5 bins according to the depth of modulation in the poststimulus window. The bins were the following: (66, –25), (–25, –37), (–37, –47), (–47, –58), (–58,– 89) % change. Here, –100% corresponds to the deepest modulation, and 0% to the absence of a change in the amplitude. **B**. P300 responses are sorted into the corresponding bins. **C**. The spatio-temporal t-test reveals clusters of significant differences between the two most extreme bins—bin 1 and bin 5. The topography of t-statistics is sampled at 500 ms (dashed line). The significant electrodes at this time point are marked with “x”.

### Spatial similarity between alpha amplitude envelope and P300 in sensor space

Consistently with the second prediction, spatial distributions of P300 and alpha amplitude modulation overlapped considerably (Spearman correlation between topographies –0.80, p-value<0.0001, Figure 4). The highest amplitude of P300 (as contrasted with sER) is localised over posterior electrodes. Similarly, the highest alpha amplitude change (also contrasted with alpha amplitude after standard stimulus) appears in the same region. The topographies were sampled at the peak of P300, which on average happened at 509±171 ms after the stimulus onset. The topography of ER (Figure 4A) was computed as the difference between the target and standard topography. The topography for alpha oscillations (Figure 4B) was computed as the ratio of amplitudes after the target and the standard stimuli. Note that the change in the alpha amplitude can be observed only through the contrast of target vs standard stimuli, since the topography of the target alpha amplitude retains prominent occipital alpha that may mask the reduction in the posterior region.

**Figure 4.**
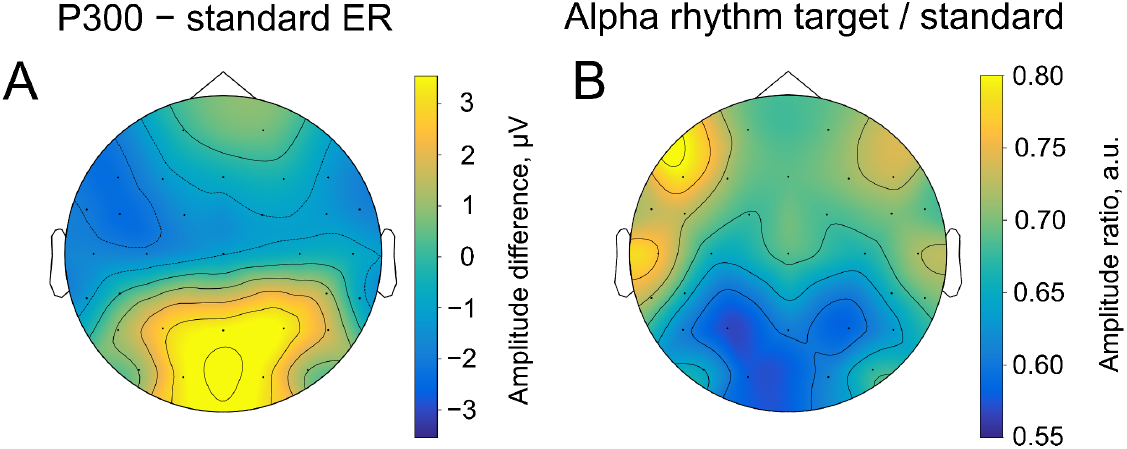
Spatial similarity of topographies of P300 (**A**) and alpha amplitude (**B**) contrasted between the target and standard stimulus. The topographies are shown at the peak amplitude of P300, which was estimated from the averaged over trials ER for each participant within the time window of 200-1000 ms poststimulus at the Pz electrode (on average 509±171 ms). For ER, the contrast was built by subtracting the sER amplitude from the P300 amplitude. For alpha amplitude, the contrast was built by dividing values of the amplitude after the target stimulus onto values after the standard stimulus.

### Spatial similarity between alpha amplitude envelope and P300 in source space

To support the sensor space spatial similarity outcome and refine the spatial overlap location between P300 and alpha amplitude changes, we performed source reconstruction. As in sensor space, we juxtaposed activations from standard and target stimuli in source space, both for P300 and the change in alpha amplitude envelope. The biggest activations for both ER and alpha amplitude were localised on the parietal midline (precuneus, posterior cingulate cortex, BA 7, 31, 23; Figures 5A,B). The location of P300 is compatible with previous studies (Tarkka et al., 1996, Tarkka et al., 1998, Faro et al., 2020) as well as with sensor space topography (Figure 4). For presentation, we outlined the overlap of dipole locations that was common for P300 and alpha amplitude change (the black line in Figures 5A,B).

**Figure 5.**
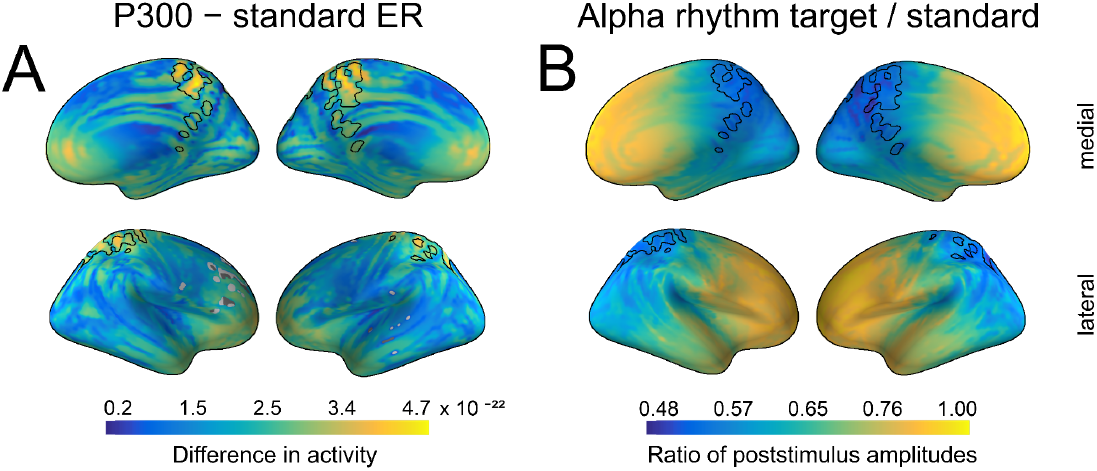
Spatial similarity between P300 and alpha amplitude in a source space. **A**. The difference between P300 and sER, after correction for multiple comparisons. The difference was estimated as the subtraction of averaged sER power from averaged P300 power in the time window of 300–700 ms. The colorbar thus indicates the difference in power. The black line outlines an overlap that is common for both P300 (top 10% of activity) and alpha amplitude (top 10% of activity). **B**. The difference in alpha amplitude envelope after standard and target stimuli with a correction for multiple comparisons (all dipole locations are significant). The difference was estimated as the target poststimulus alpha amplitude divided by the standard alpha amplitude. The poststimulus window was the same as for P300: 300–700 ms.

### The decrease in alpha amplitude and positive deflection of P300 is explained by the sign of the oscillatory mean at resting state

In support of the third prediction, the sign of BSI, which determines the sign of the oscillatory mean, should also define the alpha amplitude change with respect to the P300 polarity. That is, for oscillations with a negative mean, the attenuation of amplitude will produce an ER with a positive polarity, whereas oscillations with a positive mean will lead to an ER with a negative polarity (see also Figure 1 and Video S2). The BSIs for each participant at each electrode were estimated from a 10-min resting-state recording. The BSIs tended to be negative on average at Pz and in the nearby occipital region (Figure 6A). The distribution of BSIs at Pz was skewed towards negative values (Figure 6B), with a mean value of −0.12 and a mode of −0.85. The distribution had a trough around zero, which indicates that oscillatory activity more often was a non-zero mean.

**Figure 6.**
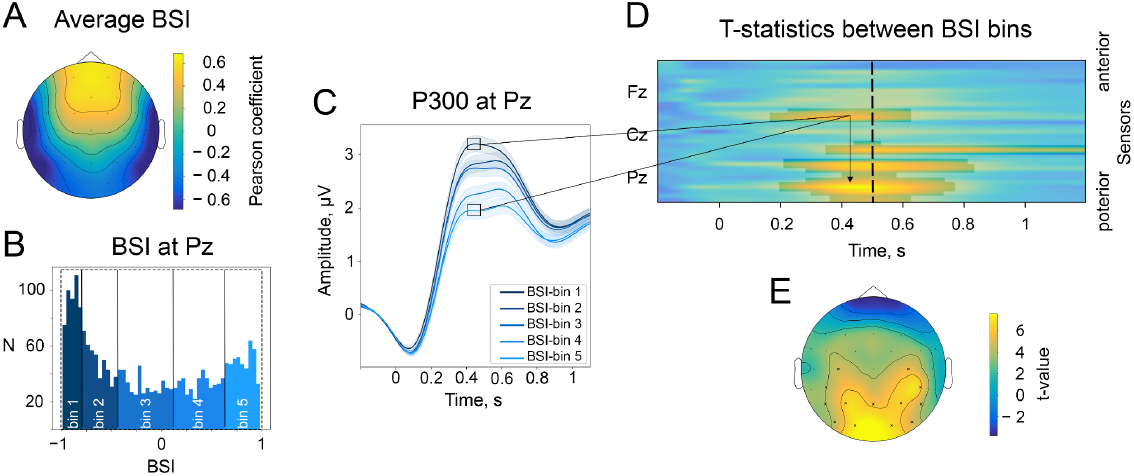
The baseline-shift index (BSI) explains the direction of ER based on the direction of alpha amplitude change. **A**. The average values of BSI at each electrode estimated from the resting-state data. Here, BSI is computed as the Pearson correlation coefficient (see Methods/The baseline-shift index). BSI serves as a proxy for the relation between ER polarity and the direction of alpha amplitude change (Nikulin et al., 2010). Here, we observe predominantly negative BSIs (and thus negative mean oscillations) at posterior sites, which indicates the inverted relation between P300 and alpha amplitude change. Indeed, in the task data, a positive deflection of P300 at posterior sites coincides with a decrease in alpha amplitude. **B**. BSIs at Pz were binned into 5 bins. The BSI bins were the following: (−0.99, −0.81), (−0.81, −0.46), (−0.46, 0.09), (0.09, 0.62), (0.62, 0.98). According to predictions of BSM, if BSI (and the oscillatory mean) was negative, then the attenuation of oscillations would lead to the upward direction of ER. **C**. P300 was binned into bins according to BSI. For bins with negative BSI, the amplitude of P300 is higher in comparison to bins with positive BSI. **D**. The evolution of the statistical difference between the amplitude of P300 in the first and fifth BSI-bins across time and space. The difference is prominent over the central and parietal regions. The cluster-based permutation test revealed significant clusters in central and parietal regions with a p-value 10^−4^. **E**. The topography of t-statistics is sampled at 500 ms (at the dashed line of the upper panel). The significant electrodes at this time point are marked with “x”.

At the sensor level, BSI computed from resting-state EEG defines the changes in P300 according to BSM (Figures 6B-E). To estimate the connection between BSI derived from the resting-state recording and P300 features, we binned BSI values into 5 bins across participants. Thus, in the first bin, there were participants with more negative BSIs (446 participants) at the particular electrode, and in the fifth bin, there were participants with positive BSIs (also 446 participants). At Pz, the BSI covaried with P300 amplitude in a way that more negative BSIs corresponded to higher amplitudes of P300 in accordance with BSM, and more positive BSIs were associated with smaller amplitudes (Figure 6C). This trend is observed in other posterior and central electrodes (Figure 6D), and we estimated significant clusters spanning electrodes FC5, C3, C4, CP5, CP6, P3, P4, P7, P8, Pz, O1, O2, PO9, PO10, and the time window of maximal P300 amplitude, approximately 300 to 700 ms (Figure 6D).

### Cognitive processes correlate with P300 and alpha amplitude modulation

Stimulus-based changes in brain signals are thought to reflect cognitive processes that are involved in the task. A simultaneous and congruent correlation of P300 and alpha rhythm to a particular cognitive score would be another evidence in favour of the relation between P300 and alpha oscillations. Moreover, if thus found, the correlation directions should correspond to the predictions according to BSM. Along with the EEG data, in the LIFE data set, a variety of cognitive tests were collected, including the Trail-making Test (TMT) A&B, Stroop test, and CERADplus neuropsychological test battery (Loeffler et al., 2015). From the cognitive tests, we extracted composite scores for attention, memory, and executive functions (Liem et al., 2017, see Methods/Cognitive tests) and tested the correlation between composite cognitive scores vs. P300 and vs. alpha amplitude modulation. The scores were available for a subset of 1549 participants (out of 2230), age range 60.03–80.01 years old. Cognitive scores correlated significantly with age (age and attention: −0.25, age and memory: −0.20, age and executive function: −0.23). Therefore, correlations between cognitive scores and electrophysiological variables were evaluated, regressing out the effect of age. To rule out the possibility of a absolute alpha power association with cognitive scores, for this analysis, we used alpha amplitude normalised change computed as 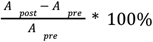, where *A*_*post*_ is at the latency of strongest amplitude decsease. Computed this way, negative alpha amplitude change would correspond to a more pronounced decrease, i.e., stronger oscillatory response.

To increase the signal-to-noise ratio of both P300 and alpha rhythm, we performed spatial filtering (see Methods/Spatial filtering, Figures 7B,C). Following this procedure, both P300 and alpha latency, but not amplitude, significantly correlated with attention scores (Figure 7A, left column). Larger latencies were related to lower attentional scores, which corresponded to a longer time-to-complete of TMT and Stroop tests and hence poorer performance. The proportion of correlation between P300 latency and attention, mediated by alpha attenuation peak latency, is 0.12. Memory scores were positively related to P300 amplitude and negatively to P300 latency (Figure 7A, middle column). The direction of correlation is such that higher memory scores, which reflected more recalled items, corresponded to a higher P300 amplitude and an earlier P300 peak. The association between alpha rhythm parameters and memory scores is not significant, but it goes in the same direction as the association for P300. Executive function (Figure 7A, right column) were related significantly to both P300 and alpha amplitude latencies. The proportion of correlation between P300 latency and attention, mediated by alpha attenuation peak latency, is 0.14. Overall, the direction of correlation is similar for P300 and alpha oscillations, as expected for BSM. Moreover, the direction of correlation is consistent across cognitive functions.

**Figure 7.**
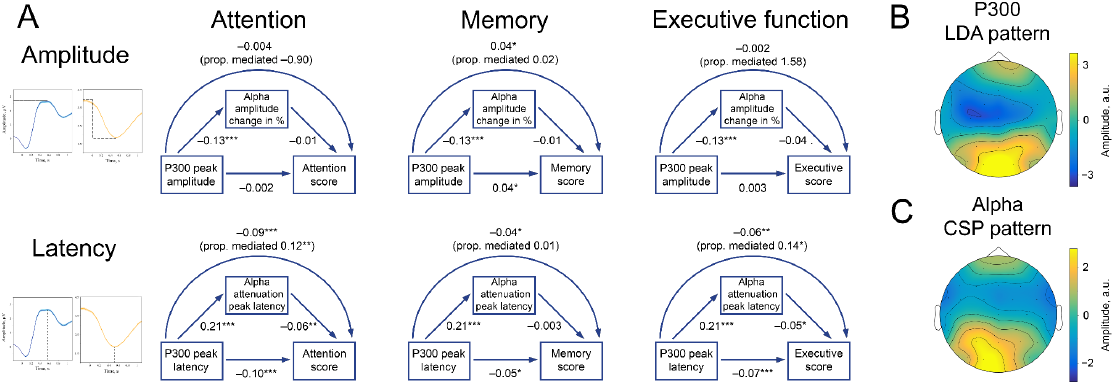
P300 and alpha oscillations showed similar correlation profiles across cognitive processes. Attention, memory, and executive function scores correlate with P300 and the alpha envelope. Attention scores were computed from TMT-A time-to-complete and Stroop-neutral time-to-complete. Memory scores were computed from the CERAD word list (combined delayed recall, recognition, and figure delayed recall). Executive function scores were computed from TMT-B time-to-complete and Stroop-incongruent time-to-complete. P300 amplitude and latency were evaluated after spatial filtering with LDA. Alpha amplitude change and latency were evaluated after spatial filtering with CSP (see Methods/Spatial filtering). Beta values were estimated with linear regression having age as a covariate variable. Sample size for this analysis is 1549.^·^p-value<0.1, * p-value<0.05, ** p-value<0.01, *** p-value<0.001. Note that the alpha amplitude change direction is such that a lower negative value would correspond to a higher decrease. B. A spatial pattern corresponding to the LDA spatial filter that was applied to obtain high signal-to-noise P300. C. A spatial pattern corresponding to the CSP filter that was applied to obtain alpha oscillations.

## Discussion

### Generation of P300 is congruent with the Baseline-shift mechanism

In the current study, we provided evidence for the hypothesis that the baseline-shift mechanism (BSM) is accountable for the generation of P300 to a certain extent. BSM for evoked response (ER) generation postulates that the modulation of oscillations with a non-zero mean leads to the generation of ER (Nikulin et al., 2007, Mazaheri et al., 2008). Here, we demonstrated the compliance of P300 generation with BSM using a large EEG data set. All the required prerequisites were confirmed: 1) the temporal courses of P300 and alpha amplitude were matching, 2) the spatial topographies of the P300 component and alpha oscillations were considerably overlapping, 3) the sign of the mean of alpha oscillations determined the direction of P300 given the decrease in alpha amplitude, 4) cognitive scores correlated in a similar way with the parameters of P300 and alpha amplitude. Therefore, P300, at least to some degree, is generated as a consequence of stimulus-triggered modulation of alpha oscillations with a non-zero mean.

The temporal correlation of P300 and alpha amplitude was negative in parietal regions. The amplitude of P300 was associated considerably with the prominence of alpha amplitude modulation, such that a smaller alpha amplitude modulation corresponded to a smaller P300 amplitude, and a larger, deeper modulation—to a larger P300. The significant cluster based on the spatio-temporal permutation test was also observed in parietal regions. Despite the fact that there is a distinct difference in P300 amplitude between participants who had a large and a small modulation, we would refrain from stating that a certain percent of P300 amplitude can be explained by alpha rhythm modulation. This conclusion cannot be definitive if we consider non-invasive recordings because spatial synchronisation within a population generating alpha rhythm greatly affects the scalp-level alpha amplitude but doesn’t affect baseline shifts (see Figure S4).

The baseline-shift index (BSI, Nikulin et al., 2010) served as a method for estimating the mean of oscillations. The topographical distribution of the BSI differed from the P300 topography. However, because BSIs were estimated from resting-state recordings, they reflect the complex neurodynamics of various alpha-frequency sources, and resting-state BSIs cannot be expected to have the same topography as P300. Yet, BSI should be non-zero in the spatial locations similar to P300 and it should have a sign compatible with the generation of P300. This is indeed what we found: BSIs in the parietal region were mostly negative, which in correspondence with the direction of P300 in relation to alpha amplitude decrease (based on BSM). Furthermore, BSI was correlated with the amplitude of P300, with a significant relation between BSI and instantaneous ER amplitude observed in centro-parietal regions in a time window of 300–600 ms after stimulus onset. The more negative BSI corresponded to higher amplitudes of P300. As posited by BSM, negative mean oscillations would generate an ER with a positive polarity.

Additionally, we tested the correlation of P300 and alpha rhythm with cognition. P300 is hypothesised to reflect attention, memory manipulation, and/or decision-making (Polich, 2007, Verleger, 2020), and previous studies showed that P300 correlated with attention (Becker et al., 1980, Nakajima et al., 2000, Lakey et al., 2011), memory (Watter et al., 2001, Braverman et al., 2003, Amin et al., 2015), and executive function (Kindermann et al., 2000, Dichter et al., 2006). Alpha rhythm has been linked to attention (Klimesch, 1999, Thut et al., 2006, Wislowska et al., 2022) and memory (Klimesch et al., 1997, Fellinger et al., 2012, van Ede, 2018, Wislowska et al., 2022). In our study, scores reflecting attention, memory, and executive function have coincidental correlations to peak amplitude and peak latency for both P300 and alpha oscillations. Namely, reduced attention and lower cognitive flexibility (Kortte et al., 2002, Douw et al., 2016) corresponded to increased peak latencies. Notably, the correlations of P300 and alpha rhythm with cognitive scores had a similar direction.

The mediation analysis showed that the modulation of alpha oscillations only partially explained the correlation between P300 and cognitive variables. This, in general, corresponds to the idea that not the whole P300 but only its fraction can be explained by the changes in the alpha amplitudes. Figure 5 shows that alpha oscillations change not only in the cortical areas where P300 is generated; therefore, we cannot expect a complete correspondence between the two processes. Moreover, since cognitive tests and EEG recordings were performed at different time points, the associations between the cognitive variables and EEG markers are expected to be rather weak and to reflect only some neuronal processes common to P300, alpha rhythm, and tasks. For these reasons, a complete mediation of one EEG variable through another EEG variable in the context of a separate cognitive assessment cannot be expected.

### Previous reports on the concurrent alpha oscillations and P300

In our review of the previous literature (presented in the Introduction and Table S1), we found a large number of studies assessing simultaneously P300 and the oscillatory dynamics in the poststimulus window. The majority of studies reveal the overlap in time windows and spatial regions of P300 and alpha amplitude decrease (see Table S1). However, not all of the studies observed a complete overlap in time courses, especially as alpha rhythm remained suppressed beyond the P300 window. Moreover, there were studies that found some discrepancies between P300 and alpha oscillations. In one study (Cooper et al., 2008), the effects of TMS were observed only in alpha oscillations. Yet, the authors admitted that, possibly, the effect on P300 was not visible because the target of TMS—the right dorsolateral prefrontal cortex—did not include the P300 sources. For other studies that failed to find the relation (Kamarajan et al., 2004, Tamura et al., 2016, Lee et al., 2017, Delval et al., 2018), we hypothesise that the evidence of the link between P300 and alpha oscillations might have been obscured due to many alpha oscillations sources being present at the same time (Rodriguez-Larios et al., 2022). Due to multiple alpha sources active at the same time, it is challenging to recover the exact alpha source that was responsible for ER generation. In particular, we observed high amplitude alpha oscillations in the occipital region (which is expected since participants were seated with their eyes closed). Moreover, the target tone presentation required participants to press the button, and as with any movement, the button press was also accompanied by oscillatory changes in the alpha (mu) frequency range (Pfurtscheller et al., 1999, Nikulin et al., 2008). In line with this assumption, we found a positive correlation between ER and alpha amplitude envelope around C3-C4 electrodes (Figure 2C) and negative ER amplitudes over the same region (Figure 4A; also see Figure S3A where the P300 time courses have negativity over central electrodes), which indicates that, possibly, there is a motor-related component of ER (Salisbury et al., 2001), with typically observed negative polarity that may have originated from a source of alpha (mu) oscillations relating to motor activity. Hence, depending on the task, there might be other changes in rhythmic activity that occlude or completely hinder the identification of oscillations that are related to the ER in question. Furthermore, none of those studies explicitly tested the compliance of the P300 generation with BSM. In our study, we extended the analysis by showing that in the same brain region, resting-state baseline shifts related to the amplitude of the non-zero mean alpha oscillations in a similar way as P300 related to stimulus-triggered alpha amplitude change. It is important to note that when assessing the interrelatedness of ER and oscillatory processes via BSM, it is necessary to evaluate all BSM predictions.

### Alternative explanations

Previously, P300 origins have been assessed according to the predictions of the additive mechanism (Fell et al., 2004, Wan et al., 2009, Herrmann et al., 2014) and the phase-reset mechanism (Fell et al., 2004, Daly et al., 2009, Wan et al., 2009). Both mechanisms have been extensively researched for different ERs, but the assessment of P300 compliance with these mechanisms is rather problematic, as is the case in general for all non-invasive measures trying to disambiguate mechanisms of ER generation (Telenczuk et al., 2010). First, the additive mechanism postulates that ER is added to the overall activity (Wood & Allison, 1981, Jervis et al., 1983, Mäkinen et al., 2005). Consequently, ER should be accompanied by an increase in total power and not only oscillatory power. However, the P300 is always accompanied by an increase in low-frequency power in the theta range, as it is its frequency range. Therefore, the predicted increase in power exclusively due to the addition of activity (Shah et al., 2004, Mazaheri et al., 2006) is impossible to disentangle based on macroscopic recordings (Telenczuk et al., 2010) and multi-unit activity is required to confirm whether an increase in power in the P300 window is of an oscillatory or non-oscillatory nature.

Moreover, in fact, BSM can also mimic the evidence for the additive mechanism, such that an ER that is generated via BSM will always be accompanied by a change in power in the low-frequency range (Figure 1). Second, the phase-reset mechanism states that ER is created when a stimulus triggers the phase alignment of oscillators in a certain frequency (Sayers et al., 1974, Makeig et al., 2002, Hanslmayr et al., 2007). Analogously, due to the frequency content of P300, there would be increased phase consistency in the theta range since ER, be it of additive or phase-reset nature, always has phase alignment. Yet, phase alignment in the poststimulus window does not contradict BSM either. We argue that the current set of predictions for the additive and the phase-reset mechanism is insufficient to confirm the generation of P300 and needs further development. As for BSM, all four BSM prerequisites (verifiable with non-invasive EEG recordings) were validated in our study, and therefore it seems reasonable to conclude that the generation of P300 is congruent with the BSM model.

The evidence presented in the current study speaks for a partial rather than an exhausting explanation of P300’s origin through BSM. The P300 is not a single ER but rather a complex. Previously (Polich, 2003, Linden, 2005), P300 was subdivided into the complex that has an earlier component—P3a—that occurs around 300 ms after stimulus onset and is more prominent in the anterior midline, and a later component—P3b—that has a latency of 500 ms and beyond and is present to a large extent in the parietal electrodes. Besides, with PCA decomposition of P300, several other components have been observed, namely slow wave and very late negativity (Steiner et al., 2014), which further indicates the complexity of the brain’s response to a target stimulus. The known and investigated mechanisms of ER generation—additive mechanism, phase-resetting mechanism, and BSM—may explain different temporal windows of one ER (Iemi et al., 2019). In our research, we found that a slow low-frequency wave of P300 may be explained by the concurrent changes in alpha amplitude via BSM. It is nonetheless feasible that a certain part of P300 might still be generated via the additive or phase-reset mechanism, although, in contrast to BSM, the prerequisites for these two types of mechanisms are hard to verify with EEG/MEG (Telenczuk et al., 2010). Moreover, determining a certain variance of P300 amplitude that can be explained by BSM is challenging when we analyse non-invasive recordings. The synchronisation within a population generating alpha rhythm affects the scalp-level alpha amplitude (see Figure S4) such that for a poorly synchronised network, the power of oscillations is severely diminished (Studenova et al., 2022). However, since the baseline shift doesn’t depend on the phase of oscillations, its amplitude is not influenced by the strength of synchronisation. Therefore, here, we did not aim to completely explain the P300 complex but to show that all four prerequisites for BSM are met for P300 generation, thus mechanistically linking P300 and alpha oscillations.

### Limitations

In our previous study (Studenova et al., 2022), using a smaller data set, we found that baseline shifts were harder to detect in the elderly population compared to the younger population. However, in the current study, due to a large sample size, we overcame difficulties related to the extraction of baseline shifts in aged participants and revealed statistically significant associations between ERs and oscillations. Essentially, the alpha amplitude decrease, triggered by the target stimulus, was particularly prominent and was substantial in the majority of participants. Only for 3% of participants, the amplitude of alpha rhythm after the target stimulus was equal to or greater than after the standard stimulus within the P300 window. In all other participants, a target stimulus evoked a pronounced attenuation of alpha oscillations. Besides, P300 in the elderly and patients with cognitive decline had smaller amplitude and longer latency (van Dinteren et al., 2014), but it never completely disappeared. In our sample, only 9% of participants had P300 amplitude smaller than sER. This in turn gave us an ample opportunity to investigate P300 and related alpha oscillations. The matter may be more complicated with other ERs that, for instance, are associated with the alpha rhythm that is generated by a smaller population of neurons and hence may be masked by other alpha rhythm sources with a higher amplitude, e.g., for auditory responses (Weisz et al., 2011).

A noteworthy limitation of the study is that EEG data was collected using only 31 channels. Spatial mixing is a substantial problem for any EEG set-up, and a smaller number of electrodes complicates the oscillatory analysis further. It was shown that with a small number of electrodes, the spatial accuracy of source reconstruction deteriorates (Liu et al., 2018, Dattola et al., 2020). In our case, in the oddball paradigm, both P300 and alpha amplitude changes were clearly detectable, and we expected that the corresponding ROIs would be rather large, and thus 31 electrode coverage would be sufficient. For other paradigms or other frequencies (like beta and gamma), the resolution of 31 channels may be insufficient since these rhythms are generated by a smaller number of neurons (Pfurtscheller et al., 1999).

### Implications of P300 and alpha rhythm relation

The detected link between P300 and alpha oscillations provides a novel avenue for the P300 interpretation, as the P300 functional role remains a subject of active discussion (Polich, 2007, Verleger, 2020). It has been suggested that P300 corresponds to the inhibition of irrelevant activity, which is needed to facilitate the processing of a stimulus or task (Polich, 2007). However, because a decrease in alpha rhythm amplitude is considered an indication of disinhibition of a particular region (Pfurtscheller et al., 1999, Jensen et al., 2010), it would follow that P300 may rather act as a correlate of activation related to the processing of the target stimulus. In previous research, alpha has been associated with attention (Foxe et al., 2011, Klimesch, 2012, Peylo et al., 2021), and working memory (Freunberger et al., 2011, de Vries et al., 2020). Attentional processes were reflected in the changes in alpha amplitude, such that it increased to suppress distractions and decreased to facilitate relevant processes (Neuper et al., 2001, van Diepen et al., 2019), while associations with working memory demonstrated inconsistent amplitude changes (Rodriguez-Larios et al., 2022). Therefore, we propose that at least partially, P300 reflects the disinhibition of regions responsible for attention and, possibly, working memory.

Here, we investigated the role of alpha oscillations in the generation of the P300 evoked response. However, our analytic pipeline may be easily applicable to any other ER that usually coincides with the modulation of any oscillations (in the form of a decrease or increase in the amplitude). The ERs suitable for testing against predictions of BSM include contingent negative variation (CNV), N400, earlier left anterior negativity (ELAN) and readiness potential (as they coincide with oscillatory changes in the alpha range, see Filipovic et al., 2001, Bastiaansen et al., 2002, Bender et al., 2004, Shibasaki & Hallett, 2006, Heimann et al., 2017). This list is not complete and may include other ERs and oscillations of higher frequencies.

In the current study, we found that the attenuation of alpha amplitude in parietal regions gives rise to the slow component of positive polarity in the P300 complex. Although sometimes analysed together, previously, P300 and alpha rhythm were not considered to represent the same neuronal process. We, on the other hand, demonstrated that alpha oscillations, at least partially, give rise to a P300 via the baseline-shift mechanism. Based on the results of our study, we suggest that general inferences about changes in P300 amplitude or latency should be derived in conjunction with changes in oscillatory dynamics. Overall, we provide a framework and evidence for the unifying mechanism responsible for the generation of evoked responses from amplitude dynamics of neuronal oscillations.

## Methods

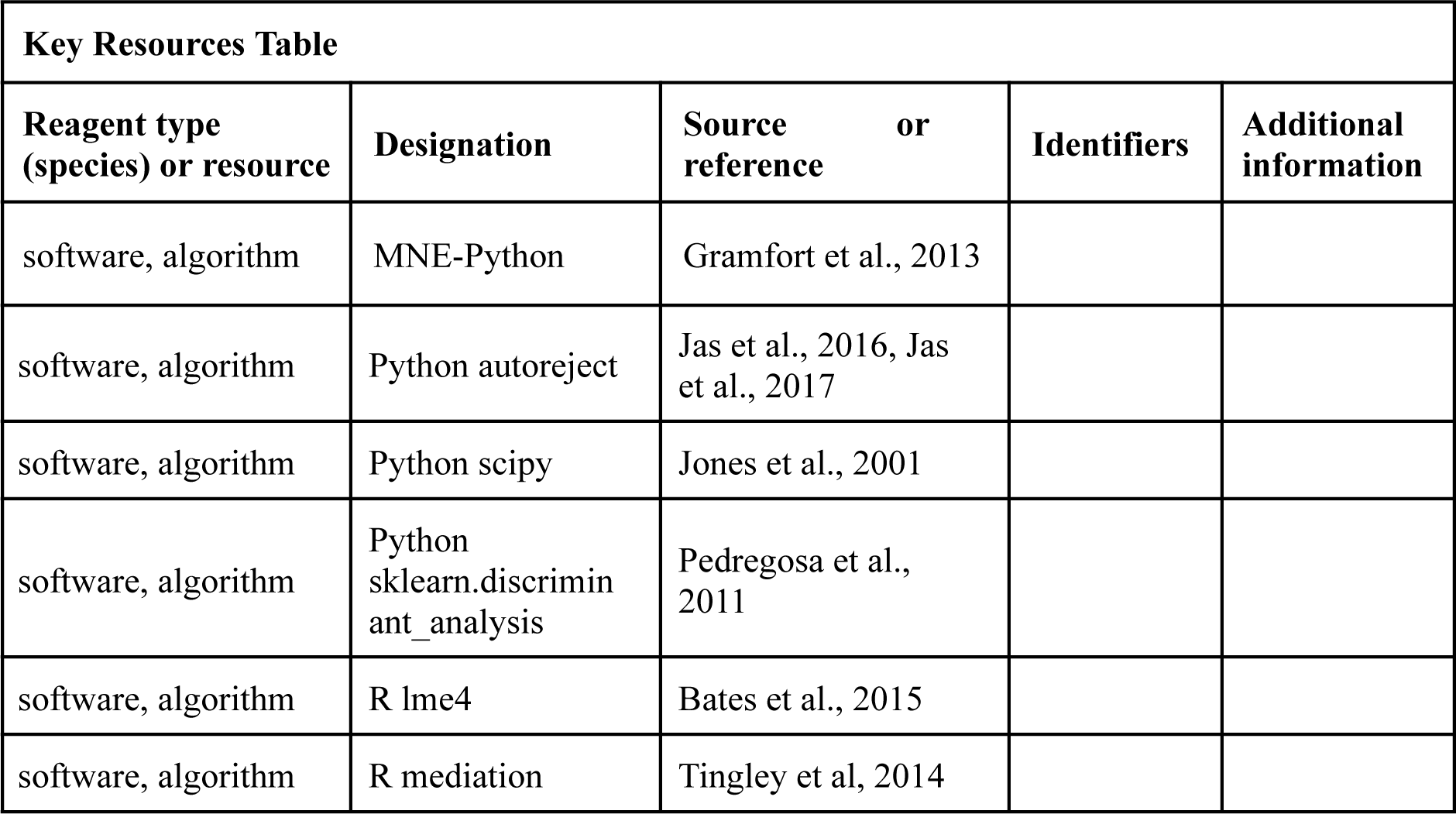

### Participants

The LIFE data set (Loeffler et al., 2015) contains data from approximately 10,000 individuals aged 40–79 years. All participants gave their written informed consent. For our study, we selected participants who took part both in resting and stimulus EEG sessions (a total of 2886 participants). From that, we had to remove 12 due to inconsistencies in stimuli coding and the mismatched header files, and 7 due to short recordings. We included all participants with no obvious neurological and psychological disorders at the moment of testing (97 participants were rejected due to medications taken at the time of data collection). We assessed the quality of the data by checking the electrode-level spectra of both resting and stimulus-based recordings. Based on the visual inspection of the quality of spectra in the low-frequency range (significant noise in more than two channels, noise in a low-frequency range of larger amplitude than the alpha peak), we rejected 539 participants (451 based on resting-state recordings, 282 based on stimulus recordings, some of them overlap). The resulting sample contained 2230 participants, aged 60–82 years old, 1152 females.

### Resting session

During the day of the recording, each participant went through three sessions: resting-state session, the oddball-novelty stimuli session, and the intensity dependence of acoustically evoked potentials session. The total time of EEG recording with preparation and follow-up did not exceed 120 minutes. The EEG resting session was recorded for a total of 20 minutes with an eyes-closed state. 31 electrodes were used for the recording, with additional electrodes for vertical and horizontal eye movements and heartbeats (40-channel QuickAmp amplifier). The electrode positions were already fastened on the cap according to the international 10–20 system. The impedances were kept under 10 kOm. The data were sampled at 1000 Hz with a low-pass filter at 280 Hz. The recording was performed with the common average reference (Jawinski et al., 2017). Before the EEG resting session, participants were situated in a reclined position, and instructed to relax and not to resist the urge to fall asleep. Based on the predictions of BSM, from the resting-state signal, we derived the association between alpha amplitude and corresponding low-frequency baseline shifts and quantified it with the baseline-shift index (BSI, see Methods/The baseline-shift index). To compute BSI, we assessed only the first 10 minutes after the beginning of the recording to decrease the possibility of the participants falling asleep.

### Oddball session

P300 was assessed by employing an acoustic oddball paradigm with three stimuli: standard, target, and novelty. A hearing test was carried out before the stimulus session to determine the hearing threshold for standard and target experimental stimuli. The hearing threshold was adjusted separately for each ear. Additionally, before the main experiment, a short test session was conducted to familiarise participants with standard and target stimuli and to make sure that they understood the instructions correctly. The main experimental session continued for 15 minutes; within that time, a total of 600 stimuli were presented in a pseudo-randomized order. At least two standard stimuli occurred between the target stimuli and no more than nine standard stimuli occurred in succession. The interstimulus interval was invariable and set to 1500 ms. The standard (more frequent) stimulus appeared with a probability of 76%.

Non-frequent stimuli, target and novelty, appeared with a 12% probability each. The standard stimulus was a sinusoidal tone with a frequency of 500 Hz, an intensity of 80 dB, and a duration of 40 ms (including a 10 ms rise and fall flanks). The target tone had the same characteristics as the standard, except for frequency, which was set to 1000 Hz for targets. The novelty stimuli were environmental or animal sounds, with an intensity of 80 dB and an average duration of 400 ms (the rise and fall flanks were selected depending on the type of tone). Participants were instructed to press the button when they heard the target stimulus. After 300 stimuli, participants had a 30-seconds break. During the break, participants were asked to change the hand used to make the response (the hand was randomly assigned at the beginning of the experimental session). In this work, we focus on the evoked response (ER) after a target stimulus, using the ER after a standard stimulus as a contrast condition.

### Cognitive tests

Along with the EEG data, the LIFE data set also includes a large number of cognitive tests (Loeffler et al., 2015). To test the correlation of P300 and alpha oscillations with cognition, we selected tests that evaluated attention, memory, and executive function—cognitive processes that, in previous research, were shown to correlate with P300 and alpha rhythm (Nakajima et al., 2000, Lakey et al., 2011, Amin et al., 2015, Dichter et al., 2006, Klimesch, 1999, Thut et al., 2006, Fellinger et al., 2012).

Attention scores were computed from the Trail-making test (TMT) and Stroop test (Liem et al., 2017). TMT is a neuropsychological test that usually includes two tasks (Reitan, 1992). The first task, also referred to as TMT-A, requires a participant to connect numbers from 1 to 25 in ascending order as quickly as possible. The second task, also referred to as TMT-B, introduces letters in addition to numbers and requires to connect both letters and numbers in an alternating fashion in ascending order. The Stroop test was performed as a computer-based colour-word interference task (Zysset et al., 2001, Scarpina et al., 2017) with two conditions—neutral and incongruent. For attentional correlates, we selected the time-to-complete metric from TMT-A and time-to-complete in the neutral condition from the Stroop test (Kynast et al., 2018, Treviño et al., 2021). In each test, not-a-number values and implausible answers were filled with the mean values of the rest of the sample. After that, both metrics were standardised with z-score (sklearn.preprocessing.StandardScaler, Pedregosa et al., 2011) and inverted (1/value). The average of two values was taken as a composite attentional score.

Memory scores were derived from the CERADplus test battery. The CERAD neuropsychological test battery was developed by the Consortium to Establish a Registry for Alzheimer’s Disease. In the LIFE data set, an authorised German version was used (www.memoryclinic.ch, Morris et al., 1988, 1989). From the CERAD panel, we selected the delayed word recall score, delayed word recognition score, and delayed figure recall score. Every score represented the number of correctly recalled or recognized words or figures divided by the total number of possible correct answers (thus the scores were in the range from 0 to 1). Deviations from the normal answers, such as a refusal to answer, were set to zero. Lastly, each score was standardised (z-transformed) and an average of three values was taken as a composite memory score.

Executive function scores were compiled using the TMT and Stroop tests (Liem et al., 2017). From TMT, we took time-to-complete in condition B, and from the Stroop test, time-to-complete in the incongruent condition. The composite executive function score of each participant was an average of standardised (z-score) inverted TMT-B time-to-complete and standardised (z-score) inverted Stroop-incongruent time-to-complete.

### Preprocessing of resting-state and stimulus-based EEG data

The preprocessing of EEG data was performed with the MNE-Python package (Gramfort et al., 2013). For each participant, we preprocessed resting-state EEG in the following way. After loading the data, we performed re-referencing to an average common reference. Then, we filtered the recording in a wide bandpass range, from 0.1 Hz to 45 Hz, with the addition of a notch filter around 50 Hz. Bad channels and bad segments were removed based on a visual inspection (Cesnaite et al., 2023) and based on markers set by the recording technician. Additionally, we visually verified the spectrum of each participant’s EEG for noisy channels. For further analysis, we exported the first ten minutes of resting-state recording. Using this time window, we computed BSI at each electrode (see Methods/The baseline-shift index).

Next, we pre-processed EEG data from a stimulus-based oddball paradigm. We applied average reference to the data and filtered it in a range from 0.1 Hz to 45 Hz, with the addition of a notch filter around 50 Hz. Continuous data were cut to trials of 1.7 s long, starting at −0.4 s before stimulus onset, and baseline corrected, with the baseline taken as −0.2,−0.05 s before stimulus onset. We applied trial rejection based on markers set by the recording technician and based on high amplitude artefacts (detected with autoreject; Jas et al., 2016, Jas et al., 2017), then we used automatic Python functions to detect eye artefacts (mne.preprocessing.create_eog_epochs) and dampened them with signal space projection (SSP, Uusitalo & Ilmoniemi, 1997), discarding one single component (conservative choice to preserve the signal of interest). To compute the single-trial ER, we low-pass filter the data using the Python scipy.signal module (Jones et al., 2001) at 3 Hz. To compute single-trial changes in alpha amplitude, we subtracted the average broadband ER from each trial, then band-pass filtered each trial around the individual alpha peak for each sensor (estimated with the spectrum obtained by Welch’s method, scipy.signal.welch) and extracted the alpha amplitude envelope with the Hilbert transform (scipy.signal.hilbert). Lastly, we averaged both the ER and the alpha amplitude envelope over trials (Figure 1).

### The baseline-shift index

The baseline-shift index (BSI, Nikulin et al., 2010) estimates a non-zero mean of oscillations based on the fluctuations in their amplitude, as proposed by the baseline-shift mechanism (BSM, Figure 8). BSM can be summarised with the equation:

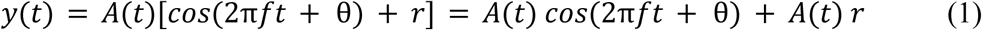

where *y*(*t*)—data from a single oscillator or a coherent population of oscillators,

**Figure 8.**
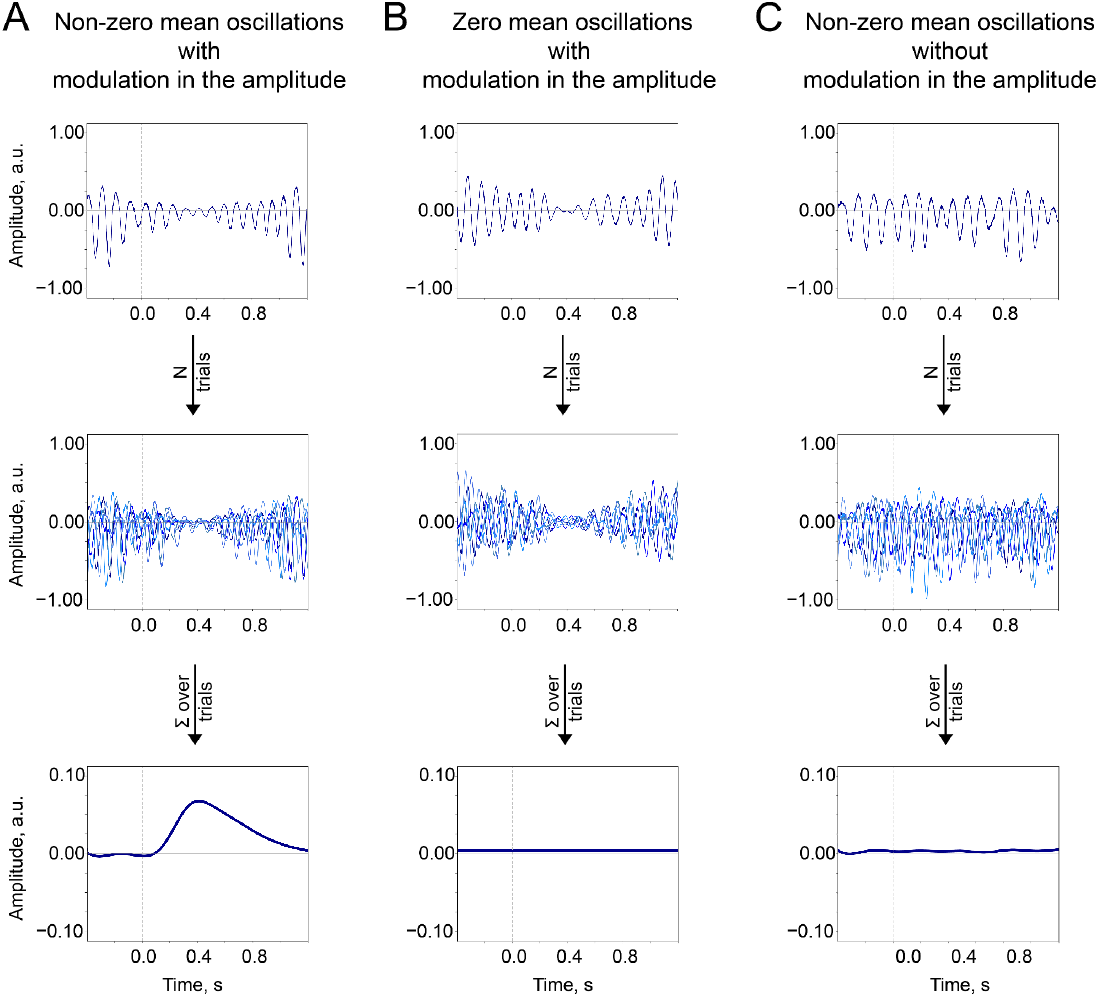
The baseline-shift mechanism (BSM) summary. Two important prerequisites of the BSM—non-zero mean *r* and amplitude modulation *A*(*t*)—should occur together so the ER would be generated. **A**. Non-zero mean oscillations when modulated in amplitude generate an ER. **B**. If oscillations have a zero mean, then no ER is generated. **C**. If oscillations have a non-zero mean but do not systematically (trial-by-trial) experience modulation, then no ER is generated.

*f*—some arbitrary frequency of oscillations in the population,

θ—some arbitrary phase,

*r*—non-zero oscillatory mean,

*A*(*t*)—amplitude modulation,

*A*(*t*) *r*—a baseline shift that accompanies oscillations.

Building on the predictions of BSM, in empirical EEG recordings, non-zero mean oscillations leave a “trace” in the low-frequency range. Therefore, the evidence of non-zero mean property for oscillations in question can be accumulated by measuring a correlation between modulation of oscillations’ amplitude (in the form of an amplitude envelope, *A*(*t*) in eq. 1) and low-frequency signal (that presumably contains baseline shifts, *A*(*t*) *r* in eq. 1). Note that the computation of BSI should be carried out using the resting-state recording to avoid contamination by stimulus effects.

A detailed description can be found in previous works (Nikulin et al., 2010, Studenova et al., 2022). In brief, firstly, we created two signals 1) by filtering broadband data in the alpha band (+−2 Hz around individual alpha peak frequency) and 2) by filtering original broadband data in the low-frequency band (low-pass at 3Hz). Filtering in the alpha band was performed with a zero-phase Butterworth filter of fourth-order, and filtering of a low-frequency signal—with a zero-phase Butterworth filter of eighth-order (scipy.signal.butter, scipy.signal.filtfilt). From filtered alpha oscillations, we derived an amplitude envelope using the Hilbert transform.

Secondly, we binned alpha amplitude into 20 bins, from the smallest to the biggest amplitude. Using the same allocation, we placed a low-frequency signal into bins as well. Amplitude values inside each bin were averaged, thus creating 20 corresponding points for alpha amplitude and low-frequency amplitude. Lastly, the relation between alpha amplitude and low-frequency amplitude was estimated as the Pearson correlation coefficient.

### The temporal and spatial similarity of alpha amplitude and P300

P300 appears in response to the target but not to the standard stimulus. Similarly, prominent alpha modulation occurs after target stimulus presentation in comparison to standard stimulus. To quantify the relation between both processes, we compared the topographical distribution of P300 and alpha amplitude dynamics in the poststimulus window around the peak of P300. We detected the peak amplitude of P300 from a filtered averaged ER in a window of 200-1000 ms at the Pz electrode. 57 participants (only 2.6% of the total number of participants) did not have an identifiable peak; those participants’ topographies were fixed at 500 ms. The topography of ER was computed as the difference between target and standard topography. The topography for alpha oscillations was computed as the ratio of amplitudes after the target and the standard stimuli. The poststimulus window for alpha amplitudes was chosen according to the ER peak latency as (*t*_*peak*_ − 50, *t*_*peak*_ + 50 ms. In the source space, the difference in evoked activations was estimated as the subtraction of averaged sER power from averaged P300 power in the time window of 300-700 ms. For the alpha amplitude envelope, the difference was estimated as the target amplitude divided by the standard amplitude in the poststimulus window 300-700 ms.

### Spatial filtering

While the averaged P300 and alpha amplitude envelope in a sensor space had distinctive similarities, not all of the participants had a high signal-to-noise ratio time course. To obtain a clearer time course estimate for each participant, we performed spatial filtering. For P300 derivation, we applied LDA (Blankertz et al., 2011, sklearn.discriminant_analysis.LinearDiscriminantAnalysis, Pedregosa et al., 2011) obtained over all participants. To achieve that, first, we computed averaged time courses of P300 and sER (ER after standard stimulus) for each participant. Second, we obtained averaged amplitude in the time window from 300 to 700 ms, thus creating two values of amplitude for each participant. Third, the LDA was trained with P300 amplitude values (matrix—participants by electrodes) and sER amplitude values (matrix—participants by electrodes) as two distinct classes. The result is the spatial filter, which is a set of weights for electrodes that maximise the difference between classes while minimising the variance inside the class, thus providing the largest discriminability between the classes. The spatial pattern was derived with the spatial filter and covariance of the cumulative data using the formula 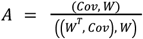, where *A*—spatial pattern, *W*—spatial filter, *Cov*—covariance of stacked data (Schaworonkow et al., 2022). Lastly, the weights derived from LDA were applied to the data of every participant, thus obtaining a single-time course of P300. From this time course, the peak amplitude and peak latency were extracted for further analysis.

For alpha oscillations, we applied a CSP spatial filter (code based on Schaworonkow et al., 2022, scipy.linalg.eig). First, we obtained covariance matrices for each participant for each condition, averaged over trials. Specifically, for each participant, we computed a covariance matrix of every alpha-filtered trial in the time window from 300 to 700 ms and then averaged trial-based matrices within the condition (Zuure et al., 2021). Second, we averaged covariance matrices to obtain a grand average over the sample of all participants. Third, using two covariance matrices of target and standard stimuli, we computed CSP filters and corresponding patterns. From those patterns, we selected the one that had the largest similarity to the P300 topography. This was the first CSP component with the largest eigenvalue. Lastly, we applied a selected filter to the data of every participant. Then the spatially filtered alpha oscillations from every trial were processed with Hilbert-transform to compute the amplitude envelope, as before (see Methods/Preprocessing of resting-state and stimulus-based EEG data). From the averaged-over-trials amplitude envelope, we derived the latency and the amplitude of the attenuation peak.

### Source reconstruction

After initial preprocessing, the stimulus-based data were filtered in the band 0.1-20 Hz and decimated to the sampling rate of 100 Hz and all trials have been concatenated for further source reconstruction. We used source localization based on the fsaverage subject (Python module mne.minimum_norm, mne-fsaverage) from FreeSurfer (Fischl, 2012). A 3-layer Boundary Element Method (BEM) model was used to compute the forward model. Source reconstruction was carried out with eLORETA (Barry et al., 2020) with the following parameters: free-orientation inverse operator (loose = 1.0), normal to the cortical surface orientation of dipoles, the regularisation parameter lambda = 0.05, and the noise covariance is the covariance of white noise signals with equal duration to the data (Idaji et al., 2022). Source spaces had 4098 candidate dipole locations per hemisphere. Reconstructed data were split into trials after reconstruction and passed through the processing pipeline as in sensor space, namely, averaging ER and computing alpha amplitude envelope (Figure 1).

### Statistical evaluation

To estimate the statistical significance for sensor-space data, if not mentioned otherwise, we used Bonferroni corrected p-values. Namely, in a sensor space, the threshold for each electrode was set as a p-value = 10^−4^/31. For source space, the threshold was chosen in a similar fashion: p-value = 10^−4^/8196.

In source space, we identified an overlap of the most prominent activity. For each dipole location on a cortical surface mesh, we computed t-statistics between the amplitude of sER and P300 in a time window from 300–700 ms. From all locations that have a significant difference, we took those that have the biggest difference in power between P300 and sER (top 10%). Similarly for alpha amplitude, we identified all significant locations, and then took 10% of those significant locations that had the biggest ratio of target to standard alpha amplitude in the poststimulus window of 300–700 ms. Then, we extracted the overlap between the two regions of interest for presentation purposes.

We ran additional statistical analysis to test the relation between P300 amplitude and the depth of alpha amplitude modulation in a sensor space. The depth of amplitude modulation was computed as 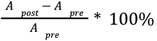. For each electrode, we binned the alpha amplitude modulation of all participants into 5 bins. Then, we used this binning to sort P300. Next, for each time point, we computed t-statistics between the amplitude of P300 at that point in the 1st and 5th bins (which corresponded to the smallest and the largest modulation respectively). To evaluate significance, we ran a cluster-based permutation test (Python mne.stats.spatio_temporal_cluster_test) with 10000 permutations and the threshold corresponding to a p-value = 10^−4^.

A separate statistical test was carried out for the relation of P300 amplitude with the BSI in sensor space. For each electrode, we binned the ERs of all participants into 5 bins according to the value of BSI at that particular electrode in the resting-state recording. For each time point, we computed t-statistics between the amplitude of P300 in the 1st and 5th BSI bins (which corresponded to the most negative BSIs and the most positive BSIs for this particular electrode). After that, we ran a cluster-based permutation test (Python mne.stats.spatio_temporal_cluster_test) with 10000 permutations and the threshold corresponding to a p-value = 10^−4^.

The correlation between cognitive scores (see Methods/Cognitive tests) and the amplitude and latency of P300 and alpha oscillations was calculated with linear regression using age as a covariate (R lme4, Bates et al., 2015). To estimate what proportion of the correlation between P300 and cognitive score is mediated by alpha oscillations, we used mediation analysis (Baron et al., 1986; R mediation, Tingley et al, 2014). First, we estimated the effect of P300 on the cognitive variable of interest (total effect, *cogscore ∼ P300+age*). Second, we computed the association between P300 and alpha oscillations (the effect on the mediator, *alpha ∼ P300*). Third, we run the full model (the effect of the mediator on the variable of interest, *cogscore ∼ P300+alpha+age*). Lastly, we estimated the proportion mediated.

## Data and code availability

Anonymised data will be made available upon request through the application procedure carried out by the LIFE-Study administration (https://www.uniklinikum-leipzig.de/einrichtungen/life). Code is available at github.com/astudenova/p300_alpha.

## Author contributions

M.L., T.H., S.C., N.M. U.H. designed the experiments and collected the data. A.A.S., D.A.E., A.V., and V.V.N wrote the manuscript. A.A.S. analysed the data. C.F. performed a code review. V.V.N. supervised the project.

## References

Amin, H.U., Malik, A.S., Kamel, N. et al. P300 correlates with learning & memory abilities and fluid intelligence. J NeuroEngineering Rehabil 12, 87 (2015). 10.1186/s12984-015-0077-6

Babiloni, C., Del Percio, C., Brancucci, A., Capotosto, P., Le Pera, D., Marzano, N., … & Rossini, P. M. (2008). Pre-stimulus alpha power affects vertex N2–P2 potentials evoked by noxious stimuli. Brain research bulletin, 75(5), 581–590.

Baron, R. M., & Kenny, D. A. (1986). The moderator–mediator variable distinction in social psychological research: Conceptual, strategic, and statistical considerations. Journal of personality and social psychology, 51(6), 1173.

Barry, R. J., Steiner, G. Z., De Blasio, F. M., Fogarty, J. S., Karamacoska, D., & MacDonald, (2020). Components in the P300: Don’t forget the Novelty P3!. Psychophysiology, 57(7), e13371.

Barutchu, A., Freestone, D. R., Innes-Brown, H., Crewther, D. P., & Crewther, S. G. (2013). Evidence for enhanced multisensory facilitation with stimulus relevance: an electrophysiological investigation. PLoS One, 8(1), e52978.

Bastiaansen, M. C., Van Berkum, J. J., & Hagoort, P. (2002). Event-related theta power increases in the human EEG during online sentence processing. Neuroscience letters, 323(1), 13–16.

Bates, D., Mächler, M., Bolker, B., Walker, S. (2015). Fitting Linear Mixed-Effects Models Using lme4. Journal of Statistical Software, 67(1), 1–48.

Becker, D. E., & Shapiro, D. (1980). Directing attention toward stimuli affects the P300 but not the orienting response. Psychophysiology, 17(4), 385–389.

Bender, S., Oelkers-Ax, R., Resch, F., & Weisbrod, M. (2004). Motor processing after movement execution as revealed by evoked and induced activity. Cognitive brain research, 21(1), 49–58.

Blankertz, B., Lemm, S., Treder, M., Haufe, S., & Müller, K. R. (2011). Single-trial analysis and classification of ERP components—a tutorial. NeuroImage, 56(2), 814–825.

Brandt, M. E., Jansen, B. H., & Carbonari, J. P. (1991). Pre-stimulus spectral EEG patterns and the visual evoked response. Electroencephalography and Clinical Neurophysiology/Evoked Potentials Section, 80(1), 16–20.

Braverman, E. R., & Blum, K. (2003). P300 (latency) event-related potential: an accurate predictor of memory impairment. Clinical Electroencephalography, 34(3), 124–139.

Cesnaite, E., Steinfath, P., Idaji, M. J., Stephani, T., Kumral, D., Haufe, S., … & Nikulin, V. V. (2023). Alterations in rhythmic and non-rhythmic resting-state EEG activity and their link to cognition in older age. NeuroImage, 268, 119810.

Chen, X., Bin, G., Daly, I., & Gao, X. (2013). Event-related desynchronization (ERD) in the alpha band during a hand mental rotation task. Neuroscience letters, 541, 238–242.

Cooper, N. R., Fitzgerald, P. B., Croft, R. J., Upton, D. J., Segrave, R. A., Daskalakis, Z. J., & Kulkarni, J. (2008). Effects of rTMS on an auditory oddball task: a pilot study of cortical plasticity and the EEG. Clinical EEG and neuroscience, 39(3), 139–143.

Daly, I., Nasuto, S. J., & Warwick, K. (2009, July). Phase resetting as a mechanism of ERP generation; evidence from the power spectrum. In Postgraduate Conference in Biomedical Engineering & Medical Physics (p. 45).

Dattola, S. et al. (2020). Effect of Sensor Density on eLORETA Source Localization Accuracy. In: Esposito, A., Faundez-Zanuy, M., Morabito, F., Pasero, E. (eds) Neural Approaches to Dynamics of Signal Exchanges. Smart Innovation, Systems and Technologies, vol 151. Springer, Singapore. 10.1007/978-981-13-8950-4_36

de Vries, I. E., Slagter, H. A., & Olivers, C. N. (2020). Oscillatory control over representational states in working memory. Trends in cognitive sciences, 24(2), 150–162.

Deiber, M. P., Ibañez, V., Missonnier, P., Rodriguez, C., & Giannakopoulos, P. (2013). Age-associated modulations of cerebral oscillatory patterns related to attention control. NeuroImage, 82, 531–546.

Deiber, M. P., Meziane, H. B., Hasler, R., Rodriguez, C., Toma, S., Ackermann, M., … & Giannakopoulos, P. (2015). Attention and working memory-related EEG markers of subtle cognitive deterioration in healthy elderly individuals. Journal of Alzheimer’s disease, 47(2), 335–349.

Delval, A., Braquet, A., Dirhoussi, N., Bayot, M., Derambure, P., Defebvre, L., … & Dujardin, K. (2018). Motor preparation of step initiation: error-related cortical oscillations. Neuroscience, 393, 12–23.

Dichter Gabriel S. PhD; van der Stelt, Odin PhD; Boch, Jane Laube BS; Belger, Aysenil PhD Relations Among Intelligence, Executive Function, and P300 Event Related Potentials in Schizophrenia, The Journal of Nervous and Mental Disease: March 2006 -Volume 194 -Issue 3 -p 179–187

Digiacomo, M. R., Marco-Pallarés, J., Flores, A. B., & Gómez, C. M. (2008). Wavelet analysis of the EEG during the neurocognitive evaluation of invalidly cued targets. Brain research, 1234, 94–103.

Dong, S., Reder, L. M., Yao, Y., Liu, Y., & Chen, F. (2015). Individual differences in working memory capacity are reflected in different ERP and EEG patterns to task difficulty. Brain research, 1616, 146–156.

Douw, L., Wakeman, D. G., Tanaka, N., Liu, H., & Stufflebeam, S. M. (2016). State-dependent variability of dynamic functional connectivity between frontoparietal and default networks relates to cognitive flexibility. Neuroscience, 339, 12–21.

Espenhahn, S., Yan, T., Beltrano, W., Kaur, S., Godfrey, K., Cortese, F., … & Harris, A. D. (2020). The effect of movie-watching on electroencephalographic responses to tactile stimulation. Neuroimage, 220, 117130.

Fabi, S., & Leuthold, H. (2017). Empathy for pain influences perceptual and motor processing: Evidence from response force, ERPs, and EEG oscillations. Social Neuroscience, 12(6), 701–716.

Fabi, S., & Leuthold, H. (2018). Racial bias in empathy: Do we process dark-and fair-colored hands in pain differently? An EEG study. Neuropsychologia, 114, 143–157.

Faro, H. K. C., Machado, D. G. D. S., Bortolotti, H., do Nascimento, P. H. D., Moioli, R. C., Elsangedy, H. M., & Fontes, E. B. (2020). Influence of judo experience on neuroelectric activity during a selective attention task. Frontiers in psychology, 10, 2838.

Fell, J., Dietl, T., Grunwald, T., Kurthen, M., Klaver, P., Trautner, P., … & Fernández, G. (2004). Neural bases of cognitive ERPs: more than phase reset. Journal of cognitive neuroscience, 16(9), 1595–1604.

Fellinger R, Gruber W, Zauner A, Freunberger R, Klimesch W (2012) Evoked traveling alpha waves predict visual-semantic categorization speed. Neuroimage 59:3379–3388.

Filipović, S. R., Jahanshahi, M., & Rothwell, J. C. (2001). Uncoupling of contingent negative variation and alpha band event-related desynchronization in a go/no-go task. Clinical Neurophysiology, 112(7), 1307–1315.

Fischl, B. (2012). FreeSurfer. Neuroimage, 62(2), 774–781.

Foxe, J. J., & Snyder, A. C. (2011). The role of alpha-band brain oscillations as a sensory suppression mechanism during selective attention. Frontiers in psychology, 2, 154.

Freunberger, R., Werkle-Bergner, M., Griesmayr, B., Lindenberger, U., & Klimesch, W. (2011). Brain oscillatory correlates of working memory constraints. Brain research, 1375, 93–102.

Gramfort, A., Luessi, M., Larson, E., Engemann, D. A., Strohmeier, D., Brodbeck, C., … & Hämäläinen, M. (2013). MEG and EEG data analysis with MNE-Python. Frontiers in neuroscience, 267.

Hanslmayr, S., Klimesch, W., Sauseng, P., Gruber, W., Doppelmayr, M., Freunberger, R., … & Birbaumer, N. (2007). Alpha phase reset contributes to the generation of ERPs. Cerebral Cortex, 17(1), 1–8.

Heimann, K. S., Uithol, S., Calbi, M., Umiltà, M. A., Guerra, M., & Gallese, V. (2017). “Cuts in Action”: A High-Density EEG Study Investigating the Neural Correlates of Different Editing Techniques in Film. Cognitive science, 41(6), 1555–1588.

Herrmann, C. S., Rach, S., Vosskuhl, J., & Strüber, D. (2014). Time–frequency analysis of event-related potentials: a brief tutorial. Brain topography, 27(4), 438–450.

Idaji, M. J., Zhang, J., Stephani, T., Nolte, G., Müller, K. R., Villringer, A., & Nikulin, V. V. (2022). Harmoni: A method for eliminating spurious interactions due to the harmonic components in neuronal data. NeuroImage, 252, 119053.

Iemi, L., Busch, N. A., Laudini, A., Haegens, S., Samaha, J., Villringer, A., & Nikulin, V. V. (2019). Multiple mechanisms link prestimulus neural oscillations to sensory responses. Elife, 8, e43620.

Ishii, R., Canuet, L., Herdman, A., Gunji, A., Iwase, M., Takahashi, H., … & Takeda, M. (2009). Cortical oscillatory power changes during auditory oddball task revealed by spatially filtered magnetoencephalography. Clinical Neurophysiology, 120(3), 497–504.

Jas, M., Engemann, D., Raimondo, F., Bekhti, Y., & Gramfort, A. (2016, June). Automated rejection and repair of bad trials in MEG/EEG. In 2016 International Workshop on Pattern Recognition in Neuroimaging (PRNI) (pp. 1–4). IEEE.

Jas, M., Engemann, D. A., Bekhti, Y., Raimondo, F., & Gramfort, A. (2017). Autoreject: Automated artifact rejection for MEG and EEG data. NeuroImage, 159, 417–429.

Jawinski, P., Kittel, J., Sander, C., Huang, J., Spada, J., Ulke, C., … & Hegerl, U. (2017). Recorded and reported sleepiness: the association between brain arousal in resting state and subjective daytime sleepiness. Sleep, 40(7).

Jensen, O., & Mazaheri, A. (2010). Shaping functional architecture by oscillatory alpha activity: Gating by inhibition. Frontiers in Human Neuroscience, 4(November), 1–8.

Jervis, B. W., Nichols, M. J., Johnson, T. E., Allen, E., & Hudson, N. R. (1983). A fundamental investigation of the composition of auditory evoked potentials. IEEE Transactions on Biomedical Engineering, (1), 43–50.

Jones, E., Oliphant, T., & Peterson, P. (2001). SciPy: Open source scientific tools for Python.

Kamarajan, C., Porjesz, B., Jones, K. A., Choi, K., Chorlian, D. B., Padmanabhapillai, A., … & Begleiter, H. (2004). The role of brain oscillations as functional correlates of cognitive systems: a study of frontal inhibitory control in alcoholism. International Journal of Psychophysiology, 51(2), 155–180.

Kamarajan, C., Porjesz, B., Jones, K., Chorlian, D., Padmanabhapillai, A., Rangaswamy, M., … & Begleiter, H. (2006). Event-related oscillations in offspring of alcoholics: neurocognitive disinhibition as a risk for alcoholism. Biological psychiatry, 59(7), 625–634.

Kao, S. C., Wang, C. H., & Hillman, C. H. (2020). Acute effects of aerobic exercise on response variability and neuroelectric indices during a serial n-back task. Brain and Cognition, 138, 105508.

Kayser, J., Tenke, C. E., Kroppmann, C. J., Alschuler, D. M., Fekri, S., Ben-David, S., … & Bruder, G. E. (2014). Auditory event-related potentials and alpha oscillations in the psychosis prodrome: neuronal generator patterns during a novelty oddball task. International Journal of Psychophysiology, 91(2), 104–120.

Kindermann, S. S., Kalayam, B., Brown, G. G., Burdick, K. E., & Alexopoulos, G. S. (2000). Executive functions and P300 latency in elderly depressed patients and control subjects. The American Journal of Geriatric Psychiatry, 8(1), 57–65.

Klimesch, W. (2012). Alpha-band oscillations, attention, and controlled access to stored information. Trends in cognitive sciences, 16(12), 606–617.

Klimesch W, Doppelmayr M, Pachinger T, Ripper B (1997) Brain oscillations and human memory: EEG correlates in the upper alpha and theta band. Neurosci Lett 238:9–12.

Klimesch W (1999) EEG alpha and theta oscillations reflect cognitive and memory performance: a review and analysis. Brain Res Brain Res Rev 29:169–195.

Kolev, V., Yordanova, J., Schürmann, M., & Başar, E. (2001). Increased frontal phase-locking of event-related alpha oscillations during task processing. International Journal of Psychophysiology, 39(2-3), 159–165.

Kortte, K. B., Horner, M. D., & Windham, W. K. (2002). The trail making test, part B: cognitive flexibility or ability to maintain set?. Applied neuropsychology, 9(2), 106–109.

Krämer, U. M., Knight, R. T., & Münte, T. F. (2011). Electrophysiological evidence for different inhibitory mechanisms when stopping or changing a planned response. Journal of cognitive neuroscience, 23(9), 2481–2493.

Kynast, J., Lampe, L., Luck, T., Frisch, S., Arelin, K., Hoffmann, K. T., … & Schroeter, M. L. (2018). White matter hyperintensities associated with small vessel disease impair social cognition beside attention and memory. Journal of Cerebral Blood Flow & Metabolism, 38(6), 996–1009.

Lakey, C. E., Berry, D. R., & Sellers, E. W. (2011). Manipulating attention via mindfulness induction improves P300-based brain–computer interface performance. Journal of neural engineering, 8(2), 025019.

Lee, J. Y., Lindquist, K. A., & Nam, C. S. (2017). Emotional granularity effects on event-related brain potentials during affective picture processing. Frontiers in Human Neuroscience, 133.

Leroy, A., Cevallos, C., Cebolla, A. M., Caharel, S., Dan, B., & Cheron, G. (2017). Short-term EEG dynamics and neural generators evoked by navigational images. PloS One, 12(6), e0178817.

Liem, F., Varoquaux, G., Kynast, J., Beyer, F., Masouleh, S. K., Huntenburg, J. M., … & Margulies, D. S. (2017). Predicting brain-age from multimodal imaging data captures cognitive impairment. Neuroimage, 148, 179–188.

Linden, D. E. (2005). The P300: where in the brain is it produced and what does it tell us?. The Neuroscientist, 11(6), 563–576.

Liu, Q., Ganzetti, M., Wenderoth, N., & Mantini, D. (2018). Detecting large-scale brain networks using EEG: Impact of electrode density, head modeling and source localization. Frontiers in neuroinformatics, 12, 4.

Liu, J., Li, J., Peng, W., Feng, M., & Luo, Y. (2019). EEG correlates of math anxiety during arithmetic problem solving: Implication for attention deficits. Neuroscience letters, 703, 191–197.

Loeffler, M., Engel, C., Ahnert, P., Alfermann, D., Arelin, K., Baber, R., … & Thiery, J. (2015). The LIFE-Adult-Study: objectives and design of a population-based cohort study with 10,000 deeply phenotyped adults in Germany. BMC public health, 15(1), 1–14.

López-Caneda, E., Rodríguez Holguín, S., Correas, Á., Carbia, C., González-Villar, A., Maestú, F., & Cadaveira, F. (2017). Binge drinking affects brain oscillations linked to motor inhibition and execution. Journal of psychopharmacology, 31(7), 873–882.

Luck, S. J. (2014). An introduction to the event-related potential technique. MIT press.

Makeig, S., Westerfield, M., Jung, T. P., Enghoff, S., Townsend, J., Courchesne, E., & Sejnowski, T. J. (2002). Dynamic brain sources of visual evoked responses. Science, 295(5555), 690–694.

Mäkinen, V., Tiitinen, H., & May, P. (2005). Auditory event-related responses are generated independently of ongoing brain activity. NeuroImage, 24(4), 961–968.

Martel, A., Arvaneh, M., Robertson, I., Smallwood, J., & Dockree, P. (2019). Distinct neural markers for intentional and unintentional task unrelated thought. bioRxiv, 705061.

Mazaheri, A., Jensen, O. (2006). Posterior α activity is not phase-reset by visual stimuli. Proceedings of the National Academy of Sciences of the United States of America, 103(8), 2948–2952. 10.1073/pnas.0505785103

Mazaheri, A., & Jensen, O. (2008). Asymmetric amplitude modulations of brain oscillations generate slow evoked responses. Journal of Neuroscience, 28(31), 7781–7787.

Michelini, G., Kitsune, V., Vainieri, I., Hosang, G. M., Brandeis, D., Asherson, P., & Kuntsi, J. (2018). Shared and disorder-specific event-related brain oscillatory markers of attentional dysfunction in ADHD and bipolar disorder. Brain topography, 31(4), 672–689.

Morris JC et al. CERAD. Part I. Clinical and neuropsychological assessment of Alzheimer’s disease. Neurology. 1989; 39(9): 1159–1165.

Morris JC et al. Consortium to establish a registry for Alzheimer’s disease (CERAD) clinical and neuropsychological assessment of Alzheimer’s disease. Psychopharmacol Bull. 1988; 24(4): 641–652.

Nakajima, Y., & Imamura, N. (2000). Relationships between attention effects and intensity effects on the cognitive N140 and P300 components of somatosensory ERPs. Clinical neurophysiology, 111(10), 1711–1718.

Neuper, C., & Pfurtscheller, G. (2001). Event-related dynamics of cortical rhythms: frequency-specific features and functional correlates. International journal of psychophysiology, 43(1), 41–58.

Nikolin, S., Tan, Y. Y., Martin, D., Moffa, A., Loo, C. K., & Boonstra, T. W. (2021). Behavioural and neurophysiological differences in working memory function of depressed patients and healthy controls. Journal of Affective Disorders, 295, 559–568.

Nikulin, V. V., Linkenkaer-Hansen, K., Nolte, G., Lemm, S., Müller, K. R., Ilmoniemi, R. J., & Curio, G. (2007). A novel mechanism for evoked responses in the human brain. European Journal of Neuroscience, 25(10), 3146–3154.

Nikulin VV, Hohlefeld FU, Jacobs AM, Curio G. Quasi-movements: a novel motor-cognitive phenomenon. Neuropsychologia. 2008 Jan 31;46(2):727–42. doi: 10.1016/j.neuropsychologia.2007.10.008. Epub 2007 Oct 22. PMID: 18035381.

Nikulin, V. V., Linkenkaer-Hansen, K., Nolte, G., & Curio, G. (2010). Non-zero mean and asymmetry of neuronal oscillations have different implications for evoked responses. Clinical Neurophysiology, 121(2), 186–193.

Paolicelli, D., Manni, A., Iaffaldano, A., Tancredi, G., Ricci, K., Gentile, E., … & Trojano, M. (2021). Magnetoencephalography and High-Density Electroencephalography Study of Acoustic Event Related Potentials in Early Stage of Multiple Sclerosis: A Pilot Study on Cognitive Impairment and Fatigue. Brain sciences, 11(4), 481.

Pedregosa, F., Varoquaux, G., Gramfort, A., Michel, V., Thirion, B., Grisel, O., … & Duchesnay, E. (2011). Scikit-learn: Machine learning in Python. the Journal of machine Learning research, 12, 2825–2830.

Peng, W., Hu, L., Zhang, Z., & Hu, Y. (2012). Causality in the association between P300 and alpha event-related desynchronization. PLoS One, 7(4), e34163.

Peylo, C., Hilla, Y., & Sauseng, P. (2021). Cause or consequence? Alpha oscillations in visuospatial attention. Trends in Neurosciences, 44(9), 705–713.

Pfurtscheller, G., Lopes Da Silva, F. H. (1999). Event-related EEG/MEG synchronization and desynchronization: Basic principles. Clinical Neurophysiology, 110(11), 1842–1857. 10.1016/S1388-2457(99)00141-8

Polich, J., & Kok, A. (1995). Cognitive and biological determinants of P300: an integrative review. Biological psychology, 41(2), 103–146.

Polich, J. (2003). Theoretical overview of P3a and P3b. Detection of change, 83–98.

Polich, J. (2007). Updating P300: an integrative theory of P3a and P3b. Clinical neurophysiology, 118(10), 2128–2148.

Popp, F., Dallmer-Zerbe, I., Philipsen, A., & Herrmann, C. S. (2019). Challenges of P300 modulation using transcranial alternating current stimulation (tACS). Frontiers in psychology, 10, 476.

Rawls, E., Miskovic, V., & Lamm, C. (2020). Delta phase reset predicts conflict-related changes in P3 amplitude and behavior. Brain research, 1730, 146662.

Reitan RM (1992). Trail Making Test: Manual for administration and scoring. Tucson, AZ: Reitan Neuropsychology Laboratory

Rodriguez-Larios, J., ElShafei, A., Wiehe, M., & Haegens, S. (2022). Visual working memory recruits two functionally distinct alpha rhythms in posterior cortex. bioRxiv.

Román-López, T. V., Caballero-Sánchez, U., Cisneros-Luna, S., Franco-Rodríguez, J. A., Méndez-Díaz, M., Prospéro-García, O., & Ruiz-Contreras, A. E. (2019). Brain electrical activity from encoding to retrieval while maintaining and manipulating information in working memory. Memory, 27(8), 1063–1078.

Salisbury, D. F., Rutherford, B., Shenton, M. E., & McCarley, R. W. (2001). Button-pressing affects P300 amplitude and scalp topography. Clinical Neurophysiology, 112(9), 1676–1684.

Sayers, B., Beagley, H. A., & Henshall, W. R. (1974). The mechanism of auditory evoked EEG responses. Nature, 247(5441), 481–483.

Scarpina, F., & Tagini, S. (2017). The stroop color and word test. Frontiers in psychology, 8, 557.

Schaworonkow N & Nikulin VV: Is sensor space analysis good enough? Spatial patterns as a tool for assessing spatial mixing of EEG/MEG rhythms. NeuroImage (2022).

Shah, A. S., Bressler, S. L., Knuth, K. H., Ding, M., Mehta, A. D., Ulbert, I., Schroeder, C. E. (2004). Neural Dynamics and the Fundamental Mechanisms of Event-related Brain Potentials. Cerebral Cortex, 14(5), 476–483. 10.1093/cercor/bhh009

Shibasaki, H., & Hallett, M. (2006). What is the Bereitschaftspotential?. Clinical neurophysiology, 117(11), 2341–2356.

Shou, G., & Ding, L. (2015). Detection of EEG spatial–spectral–temporal signatures of errors: a comparative study of ICA-based and channel-based methods. Brain topography, 28(1), 47–61.

Steiner, G. Z., Barry, R. J., & Gonsalvez, C. J. (2014). Nontarget-to-nontarget interval determines the nontarget P300 in an auditory equiprobable Go/NoGo task. International journal of psychophysiology, 92(3), 113–121.

Studenova, A. A., Villringer, A., & Nikulin, V. V. (2022). Non-zero mean alpha oscillations revealed with computational model and empirical data. PLoS computational biology, 18(7), e1010272.

Tamura, K., Mizuba, T., & Iramina, K. (2016). Hearing subject ’ s own name induces the late positive component of event-related potential and beta power suppression. Brain Research, 1635, 130–142.

Tang, D., Hu, L., Lei, Y., Li, H., & Chen, A. (2015). Frontal and occipital-parietal alpha oscillations distinguish between stimulus conflict and response conflict. Frontiers in Human Neuroscience, 9, 433.

Tarkka, I. M., & Stokic, D. S. (1998). Source localization of P300 from oddball, single stimulus, and omitted-stimulus paradigms. Brain Topography, 11(2), 141–151.

Tarkka, I. M., Micheloyannis, S., & Stokić, D. S. (1996). Generators for human P300 elicited by somatosensory stimuli using multiple dipole source analysis. Neuroscience, 75(1), 275–287.

Telenczuk, B., Nikulin, V. V., & Curio, G. (2010). Role of neuronal synchrony in the generation of evoked EEG/MEG responses. Journal of neurophysiology, 104(6), 3557–3567.

Tingley, D., Yamamoto, T., Hirose, K., Keele, L., Imai, K. (2014). mediation: R Package for Causal Mediation Analysis. Journal of Statistical Software, 59(5), 1–38

Thut G, Nietzel A, Brandt SA, Pascual-Leone A (2006) a-Band electroencephalographic activity over occipital cortex indexes visuospatial attention bias and predicts visual target detection. J Neurosci 26:9494–9502.

Treviño, M., Zhu, X., Lu, Y. Y., Scheuer, L. S., Passell, E., Huang, G. C., … & Horowitz, T. S. (2021). How do we measure attention? Using factor analysis to establish construct validity of neuropsychological tests. Cognitive Research: Principles and Implications, 6(1), 1–26.

Uusitalo, M. A., Ilmoniemi, R.J. “Signal-space projection method for separating MEG or EEG into components.” Medical and biological engineering and computing 35.2 (1997): 135–140.

Vallat, R. (2018). Pingouin: statistics in Python. Journal of Open Source Software, 3(31), 1026, 10.21105/joss.01026

van Diepen, R. M., Foxe, J. J., Mazaheri, A. (2019). The functional role of alpha-band activity in attentional processing: the current zeitgeist and future outlook. Current Opinion in Psychology, 29, 229–238

van Dinteren, R., Arns, M., Jongsma, M. L., & Kessels, R. P. (2014). P300 development across the lifespan: a systematic review and meta-analysis. PloS one, 9(2), e87347.

van Ede, F. (2018). Mnemonic and attentional roles for states of attenuated alpha oscillations in perceptual working memory: a review. European Journal of Neuroscience, 48(7), 2509–2515.

Verleger, R. (2020). Effects of relevance and response frequency on P3b amplitudes: Review of findings and comparison of hypotheses about the process reflected by P3b. Psychophysiology, 57(7), e13542.

Vilà-Balló, A., François, C., Cucurell, D., Miró, J., Falip, M., Juncadella, M., & Rodríguez-Fornells, A. (2017). Auditory target and novelty processing in patients with unilateral hippocampal sclerosis: A current-source density study. Scientific reports, 7(1), 1–10.

Wan, B., An, X., Ming, D., Qi, H., Hu, Y., & Luk, K. D. K. (2009, May). Phase resetting and evoked activity contribute to the genesis of P300 signal in BCI system. In 2009 IEEE International Conference on Computational Intelligence for Measurement Systems and Applications (pp. 62–65). IEEE.

Watter, S., Geffen, G. M., & Geffen, L. B. (2001). The n-back as a dual-task: P300 morphology under divided attention. Psychophysiology, 38(6), 998–1003.

Weisz, N., Hartmann, T., Müller, N., Lorenz, I., & Obleser, J. (2011). Alpha rhythms in audition: cognitive and clinical perspectives. Frontiers in psychology, 2, 73.

Wislowska, M., Klimesch, W., Jensen, O., Blume, C.; Schabus, M. (2022). Sleep-Specific Processing of Auditory Stimuli Is Reflected by Alpha and Sigma Oscillations. The Journal of Neuroscience, 42 (23), 4711–4724. 10.1523/jneurosci.1889-21.2022

Wood, C. C., & Allison, T. (1981). Interpretation of evoked potentials: a neurophysiological perspective. Canadian Journal of Psychology/Revue canadienne de psychologie, 35(2), 113.

Wu, S., Hitchman, G., Tan, J., Zhao, Y., Tang, D., Wang, L., & Chen, A. (2015). The neural dynamic mechanisms of asymmetric switch costs in a combined Stroop-task-switching paradigm. Scientific reports, 5(1), 1–11.

Yordanova, J., Kolev, V., & Polich, J. (2001). P300 and alpha event-related desynchronization (ERD). Psychophysiology, 38(1), 143–152.

Yu, L., Wang, X., Lyu, Y., Ding, L., Jia, J., Tong, S., & Guo, X. (2020). Electrophysiological evidences for the rotational uncertainty effect in the hand mental rotation: an ERP and ERS/ERD study. Neuroscience, 432, 205–215.

Zarka, D., Cevallos, C., Petieau, M., Hoellinger, T., Dan, B., & Cheron, G. (2014). Neural rhythmic symphony of human walking observation: Upside-down and Uncoordinated condition on cortical theta, alpha, beta and gamma oscillations. Frontiers in systems neuroscience, 8, 169.

Zhang, Y., Liu, B., & Gao, X. (2020). Spatiotemporal dynamics of working memory under the influence of emotions based on EEG. Journal of Neural Engineering, 17(2), 026039.

Zuure, M. B., & Cohen, M. X. (2021). Narrowband multivariate source separation for semi-blind discovery of experiment contrasts. Journal of Neuroscience Methods, 350, 109063.

Zysset, S., Müller, K., Lohmann, G., von Cramon, DY, 2001. Color-word matching Stroop task: separating interference and response conflict. NeuroImage 13, 29–36.

