## Supplementary material for "Event-related modulation of alpha rhythm explains the auditory P300 evoked response in EEG"

Table S1. Overview of the previous findings concerning stimulus-related alpha amplitude decrease in an oddball paradigm and similar experiments.

The search for this short review was completed via Google Scholar on 21-09-2021 using keywords: “evoked response”, “erp”, “erf”, “erd”, “ers”, and on 03-10-2021 using keywords: “P300”, “erd”, “ers”, “eeg”, “meg”. We picked only research or review papers written in the English language.

| **Reference** | **Experimental paradigm** | **P300 localisation** | **P300 latency** | **Alpha amplitude localization** | **Alpha amplitude latency** |
|---|---|---|---|---|---|
| Kolev et al., 2001 | passive auditory oddball task | Pz | 300–500 ms | Pz | 300–800 ms |
| Yordanova et al., 2001 | active and passive auditory oddball task | Fz, Cz, Pz | average 347 ms | Fz, Cz, Pz | average 680 ms |
| Kamarajan et al., 2004 | Go-Nogo task | Pz | 400–800 ms | no alpha | - |
| Kamarajan et al., 2006 | Go-Nogo task | Cz | 300–600 ms | Cz | 300–1000 ms |
| Cooper et al., 2008 | passive auditory oddball task | Cz | 280–450 ms | all over the cortex | 150–650 ms |
| Digiacomo et al., 2008 | visual stimuli, validly and invalidly cued | Fz, FCz, Cz, Pz | 300–500 ms | Fz, FCz, Cz, Pz, P3, O2 | 300–800 ms |
| Ishii et al., 2009 | auditory oddball task | central, non-dipolar pattern (MEG) | 300–400 ms | increase in the amplitude over prefrontal cortex (bilateral superior frontal gyrus), decrease in amplitude over sensorimotor cortex | 200–600 ms |

| | | | | | |
|---|---|---|---|---|---|
| | | | | (bilateral postcentral gyrus) | |
| Krämer et al., 2011 | Eriksen flanker task | Fz, Cz, Pz | 300–700 ms | C3, C4 | 200–800 ms |
| Peng et al., 2012 | auditory oddball task paired with somatosensory stimuli | Pz | 300–800 ms | parietal region | 300–600 ms |
| Barutchu et al., 2013 | audiovisual discrimination task | Oz, O1, O2 | 250–500 ms | O1, O2 | 200–800 ms |
| Chen et al., 2013 | hand mental rotation task | Cz, Pz | 300–600 ms | CP3 | 300–600 ms |
| Deiber et al., 2013 | visual attention network test | Pz | 300–600 ms | P3, P4, Pz | 300–2000 ms |
| Kayser et al., 2013 | auditory novelty oddball task | Pz | 250–700 ms | CP2 | 400–800 ms |
| Shou et al., 2015 | Stroop-task-switching paradigm | Pz | 300–500 ms | Pz | 300–500 ms |
| Zarka et al., 2014 | video stimuli with animation | O1, O2, P3, P4 | 250–500 ms | O1, O2, P3, P4 | starts at 300 ms |
| Deiber et al., 2015 | 2-back working memory task | Cz, Pz | 400–1000 ms | O2 | 400–800 ms |
| Dong et al., 2015 | Modified Digit Span task | Pz | 300–800 ms | parietal region (P3, Pz, P4) | 350–1100 ms |
| Tang et al., 2015 | color-word flanker task | centro-parietal region | 300–600 ms | occipito-parietal region | 300–600 ms |
| Wu et al., 2015 | Stroop-task-switching paradigm | parieto-central region | 300–800 ms | fronto-central region | 300–1000 ms |
| Tamura et al., 2016 | auditory presentation of subject's own name and other names | midline | 300–700 ms | C4 | 200–600 ms |

| | | | | | |
|---|---|---|---|---|---|
| Fabi et al., 2017 | categorization of pictures with painful and non-painful stimuli | Pz | 400–800 ms | sensorimotor cortex | 300–1000 ms |
| Lee et al., 2017 | affective images presentation | posterior region | 300–400 ms | posterior region | 100–500 ms |
| Leroy et al., 2017 | visual task involving presentation of 2D and 3D images | O2 | 300–600 ms | O2 | 100–500 ms |
| López-Caneda et al., 2017 | Go-Nogo task | Fz, Cz, Pz | 300–700 ms | Pz | 400–600 ms |
| Vilà-Balló et al., 2017 | auditory novelty oddball task | Pz | 300–800 ms | Pz | 300–800 ms |
| Delval et al., 2018 | visual stimuli following gate initiation | Pz | 400–700 ms | Cz | no change in the amplitude |
| Fabi et al., 2018 | categorization of pictures with painful and non-painful stimuli | Pz | 400–700 ms | FC1, FC2, C1, C2, C3, C4, CP1, CP2 | 300–1000 ms |
| Michelini et al., 2018 | four-choice reaction time task | Pz | 300–500 ms | parieto-occipital region | 300–1000 ms |
| Liu et al., 2019 | arithmetic problem-solving | O1 | 200–700 ms | PO4, PO8 | 200–800 ms |
| Martel et al., 2019 | Go-Nogo task | centro-parietal region | 268–388 ms | all channels | 200–600 ms |
| Román-López et al., 2019 | delayed-match to sample task | midline | 300–700 ms | midline | 200–1000 ms |
| Faro et al., 2020 | Stroop colour-word test | precuneus | 250–400 ms | precuneus | 200–1000 ms |

| | | | | | |
|---|---|---|---|---|---|
| Espenhahn et al., 2020 | video viewing while passive tactile stimulation | somatosensory cortex | 270–340 ms | somatosensory cortex | 200–400 ms |
| Kao et al., 2020 | n-back working memory task | Cz, CPz, Pz | 300–800 ms | Fz, FCz | 200–900 ms |
| Yu et al., 2020 | hand mental rotation task | Pz | 300–600 ms | Pz | 200–600 ms |
| Zhang et al., 2020 | change detection paradigm | occipital region | 300–600 ms | occipital region | 300–700 ms |
| Nikolin et al., 2021 | n-back working memory task with target stimuli and distractor cues | Fz | 300–500 ms | Fz | 300–700 ms |
| Paolicelli et al., 2021 | auditory oddball task | Pz | 280.07–342.67 ms | parietal midline | 200–800 ms |

https://osf.io/b7ke3

Video S2. The demonstration of the baseline-shift mechanism for negative and positive non-zero mean oscillations that experience a stimulus-triggered increase or decrease in the amplitude.

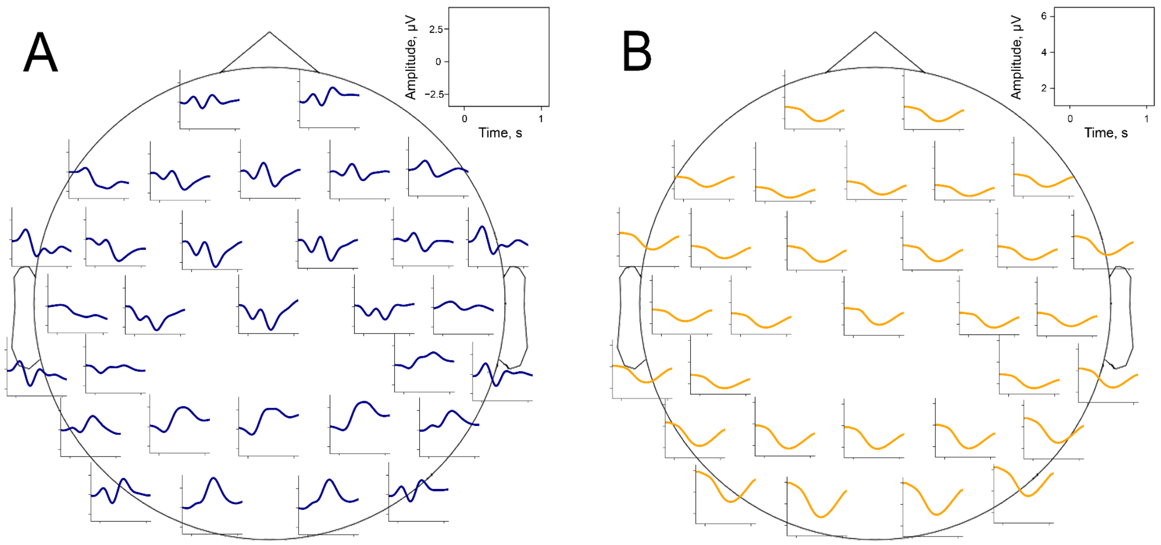


Figure S3. Time-space evolution of P300 and alpha amplitude envelope. **A**. P300 grand average over participants. **B**. Alpha amplitude grand average over participants.

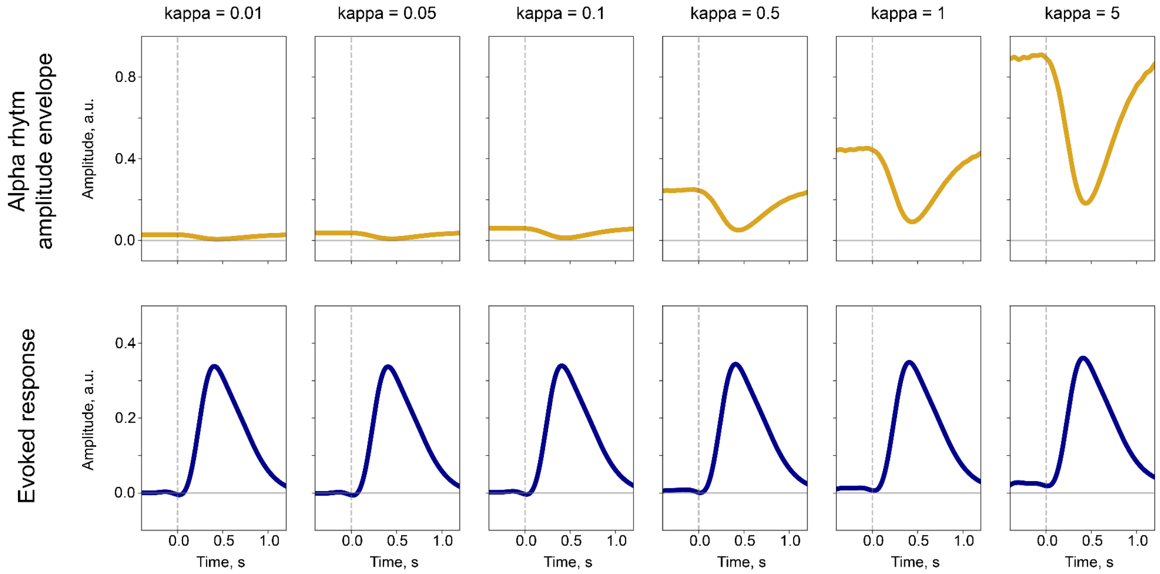


Figure S4. The synchronisation in the population of neurons generating alpha rhythm affects the amplitude of the alpha rhythm but not the evoked response. Similarly as in Studenova et al. (2022), we performed simulations with evoked response and alpha rhythm resembling P300 and alpha amplitude envelope obtained from real recordings. The synchronisation strength was manipulated by the starting distribution of phases. The kappa parameter characterises the peakiness of phase distribution—with kappa = 5 the distribution has a peak and with kappa = 0.01 the distribution is close to uniform. In the real EEG data, the synchronisation of the underlying population is unknown.

Therefore, a direct correspondence between the amplitude of the evoked response and the amplitude envelope of the alpha rhythm can be only approximately evaluated (for instance, with the baseline-shift index, BSI).

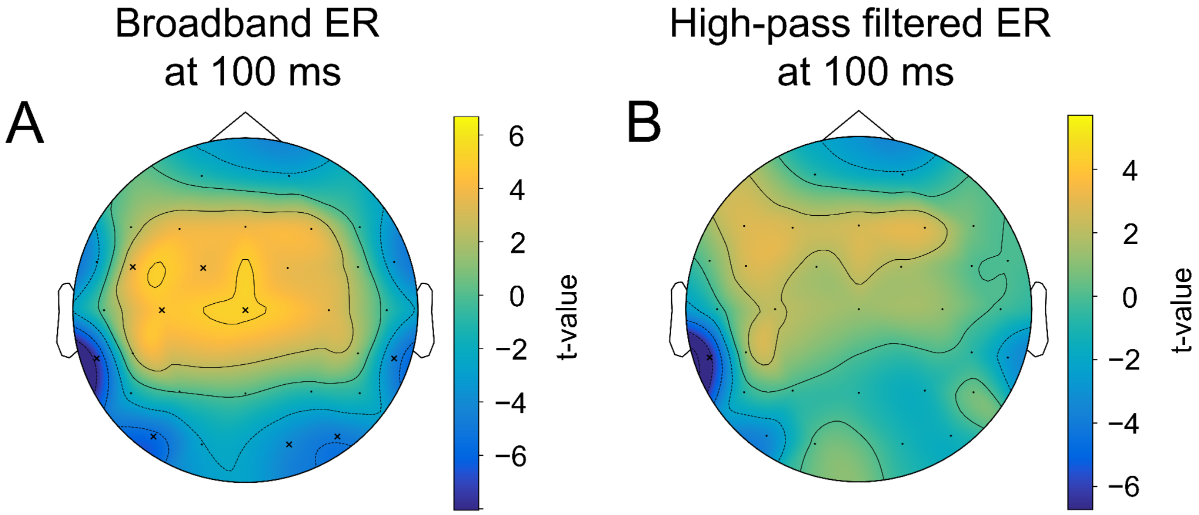


Figure S5. The difference in the strength of alpha amplitude modulation correlates with the difference in early ER, but only for broadband data. A. The spatio-temporal t-test between the two most extreme bins—bin 1 and bin 5 of alpha amplitude change (see main text, Figure 3A)—reveals a cluster of a significant difference at 100 ms (corresponding to N100) for the broadband ER (0.1–45 Hz). The significant electrodes at this time point are marked with an "x". B. The same test but for high-pass filtered ER (4–45 Hz). In this case, a large cluster of significant differences disappears. It means that the difference is driven by a low-frequency component, which presumably is a baseline shift created by an emerging decrease in alpha rhythm amplitude.
